# Cell Type-Specific Hormonal Signaling Configures Hypothalamic Circuits for Parenting

**DOI:** 10.64898/2025.12.11.693766

**Authors:** Brandon L. Logeman, Patricia M. Horvath, Mustafa Talay, Vikrant Kapoor, Harris S. Kaplan, Catherine Dulac

## Abstract

Parenting behavior emerges from hormonally sensitive circuits, but how distinct circuit components are affected by, and contribute to, sex and state dependent changes in infant caregiving remains unclear. Using cell type-specific approaches, we characterized two circuit nodes that are differentially configured by hormones to modulate infant-evoked behavior. An excitatory neuronal population in the anteroventral periventricular nucleus is only active in lactating mothers, increases virgin female caregiving when artificially stimulated and confers oxytocin sensitivity in mothers through a prolactin–STAT5b pathway. These neurons function upstream of another preoptic area population involved in male and female parenting, thus boosting caregiving by mothers. By contrast, androgen signaling in the latter preoptic population reshapes their intrinsic properties to promote pup-directed aggression, revealing cell type-specific tuning of social behavior circuits.

## Main Text

Mammals invest considerable resources to nurture offspring and undergo extensive reorganization of behavioral and physiological priorities to maximize the survival and well-being of caregiver and infants (*1–3*). While parenting is essential for survival across generations, the lack of immediate benefits for the caregiver suggests that this behavior is, at least in part, driven by evolutionarily shaped, genetically pre-programmed neural circuits. Depending on the species, parental care is provided by one or both parents, sometimes the larger social group, and lasts for a few days, months or years (*4*, *5*). Hormonal and environmental factors have been identified as key modulators of infant directed behaviors (*3*), underscoring the complexity of the neural control of parenting according to the animal internal state within a given environmental context. How systemic factors such as hormones lead to highly specific and stereotyped changes in behavior depending on sex or mating status is poorly understood. Recent efforts have aimed to uncover the specific cell types and circuits that underlie parental care (*1*, *3*, *6–22*), but whether and how specific neuronal populations within a circuit are modulated to achieve the observed state and context-dependent infant-evoked behavior has not been addressed.

Laboratory mice display a range of parental care towards infants according to their sex and physiological state. Lactating mothers display highly robust and reliable parental behavior, whereas fathers and virgin females show weaker and more variable caregiving and virgin males are typically aggressive or neglectful towards pups (*1*, *3*). What factors and mechanisms determine differences in pup-directed behavior? Decades of work on the regulation of parental care identified hormones as key determinants of adult-infant interactions. Parabiosis experiments in which virgin females were exposed to the blood plasma of lactating mothers were sufficient to induce the retrieval of infants to the virgin’s nest (*23*, *24*). Accordingly, a large number of hormones and neuropeptides including prolactin, estrogen, and oxytocin have been shown to play a key role in inducing maternal-like behaviors in otherwise non-parental animals (*3*). Conversely, in males, removal of circulating androgens through castration decreases aggression towards infants (*25*, *26*). Although these circulating hormones affect all tissues, key brain areas involved in the control of parental care express high levels of hormone receptors suggesting specific hormonal modulation of neural populations and circuits (*27–30*).

The medial preoptic area of the hypothalamus (MPOA) is essential for parental care across vertebrate species. MPOA lesions result in impaired parental care in birds and mammals (*31–41*), whereas increased MPOA neural activity in fish, frogs, and mammals is associated with parental display (*42–47*). Within the MPOA, recent studies have sought to characterize the specific circuits and cell types driving infant-evoked behaviors. MPOA neurons expressing the neuropeptide Galanin (*Gal*) were identified based on their activity during parenting and their essential role in the display of parenting behavior in both males and females (*7*, *8*). Recordings from brain slices reveal an excitatory role of the hormone prolactin on MPOA^Gal+^ neurons, suggesting their direct hormonal modulation (*19*). The specific subset of *Gal* expressing neurons activated during parenting was narrowed down to a single transcriptomic cluster co-expressing *Gal* and calcitonin receptor (*Calcr*), hereby referred to as MPOA^Gal+CalcR+^ (*9*). MPOA^Gal+CalcR+^ neurons express the estrogen (*Esr1*), androgen (*Ar*), and prolactin (*Prlr*) receptors, suggesting the integration of multiple hormonal cues by this neuronal type to direct parenting behavior (*9*). By contrast, most studies so far have examined hormonal sensitivity in broad, heterogeneous neuronal populations. For example, MPOA neuronal populations initially characterized by the expression of either *Esr1-, Calcr-*, or bombesin-like receptor 3 (*Brs3*), can now be defined as encompassing multiple and largely overlapping cell types, that influence pup retrieval but also other social and non-social functions such as aggression and temperature regulation (*15*, *22*, *48–50*).

Additional parenting-related populations reside in the anteroventral periventricular nucleus (AvPe) of the preoptic area. Tyrosine hydroxylase (TH)-expressing AvPe neurons were shown to influence maternal, but not paternal, behavior, in part through their monosynaptic projections onto *Oxt* expressing neurons in the paraventricular nucleus of the hypothalamus (PVH) and thus modulating state-dependent hormonal release to stimulate parenting behavior (*51*). The AvPe also contains an excitatory subset of the broad Brs3-expressing population in the preoptic area, distinguished by the co-expression of *Brs3* and vesicular glutamate transporter 2 (*Vglut2*). These AvPe^Brs3+Vglut2+^ neurons express high levels *Prlr*, and are *Fos*-positive in lactating mothers, but not virgin females or mated-males engaged in parenting, thus representing a prominent cell type displaying sex- and state-specific activity during parenting (*9*).

Multiple cell types have also been implicated in infant-directed aggression. The rhomboid nucleus of the bed nuclei of the stria terminalis (BSTrh) shows increased activity in infanticidal animals, with *Esr1* positive neurons driving attack toward infants (*16*, *52*). Likewise, neurons located in the perifornical area (PeFA) that express the neuropeptide urocortin 3 (Ucn3) and *Vglut2* have been implicated in infant directed aggression typically observed in virgin males (*14*, *53*).

The precise identification of specific cell types and associated neural circuits underlying the control of infant-evoked behavior offers a unique opportunity to uncover how these populations undergo sex- and state-dependent activity regulation. Internal states and external environments can affect behavior through extrinsic, network-driven changes in neural activity as well as cell-intrinsic modulation of the corresponding circuits and cell types (*54–59*). Here we hypothesize that cell-intrinsic changes in transcriptional and biophysical properties of key regulatory hubs of parenting behavior are instrumental in driving observed differences in neural activity and behavior. Using single-cell genomics and cell type-specific genetic manipulations, we uncovered sex- and physiological state-dependent remodeling of cell types controlling parental care. We identified unique targets of hormonal signaling that drive changes in cell-intrinsic properties and determine the nature of adult-infant interactions. Overall, these findings demonstrate how internal states influence neural activity in a cell type-specific manner to configure neural circuits and modulate behavior.

### A maternal-specific neural circuit

To uncover cell-intrinsic mechanisms underlying the sex- and state-dependent modulation of parenting, we focused on three cell types previously associated with adult-infant interactions: AvPe^Brs3+Vglut2+^ neurons activated during mother-specific parenting (*9*), MPOA^Gal+CalcR+^ neurons driving parenting across all parental animals (*7–9*), and PeFA^Ucn3+Vglut2+^ neurons involved in male and female infant-directed aggression (*14*, *60*) (Fig. 1 A-C). Among parental mice, mothers display the most robust infant-evoked responses compared to virgin females and fathers. To gain insights into the unique, cell type-specific, molecular features of parental regulation in lactating mothers (*34*, *38*) we sought to better characterize AvPe^Brs3+Vglut2+^ neurons, an excitatory population restricted to the AvPe that was previously found to be exclusively activated in lactating mothers but not in parental virgin females nor fathers based on immediate-early gene (IEG) expression analysis (*9*). AvPe^Brs3+Vglut2+^ neurons are part of a broader population of Brs3+ neurons distributed throughout the preoptic area that comprises cell types such as MPOA*^Gal+CalcR+^*, AvPe*^Brs3+Vglut2+^*, BNST*^Crh+/Vgf+^*, and BNST/MPA*^Aro+/Cplx3+^*, all of which are implicated in diverse social behaviors such as mating and aggression (*9*) and active during parenting in animals of various physiological states (*9*, *49*), highlighting the need for more precise cell type-specific analysis.

**Fig. 1.**
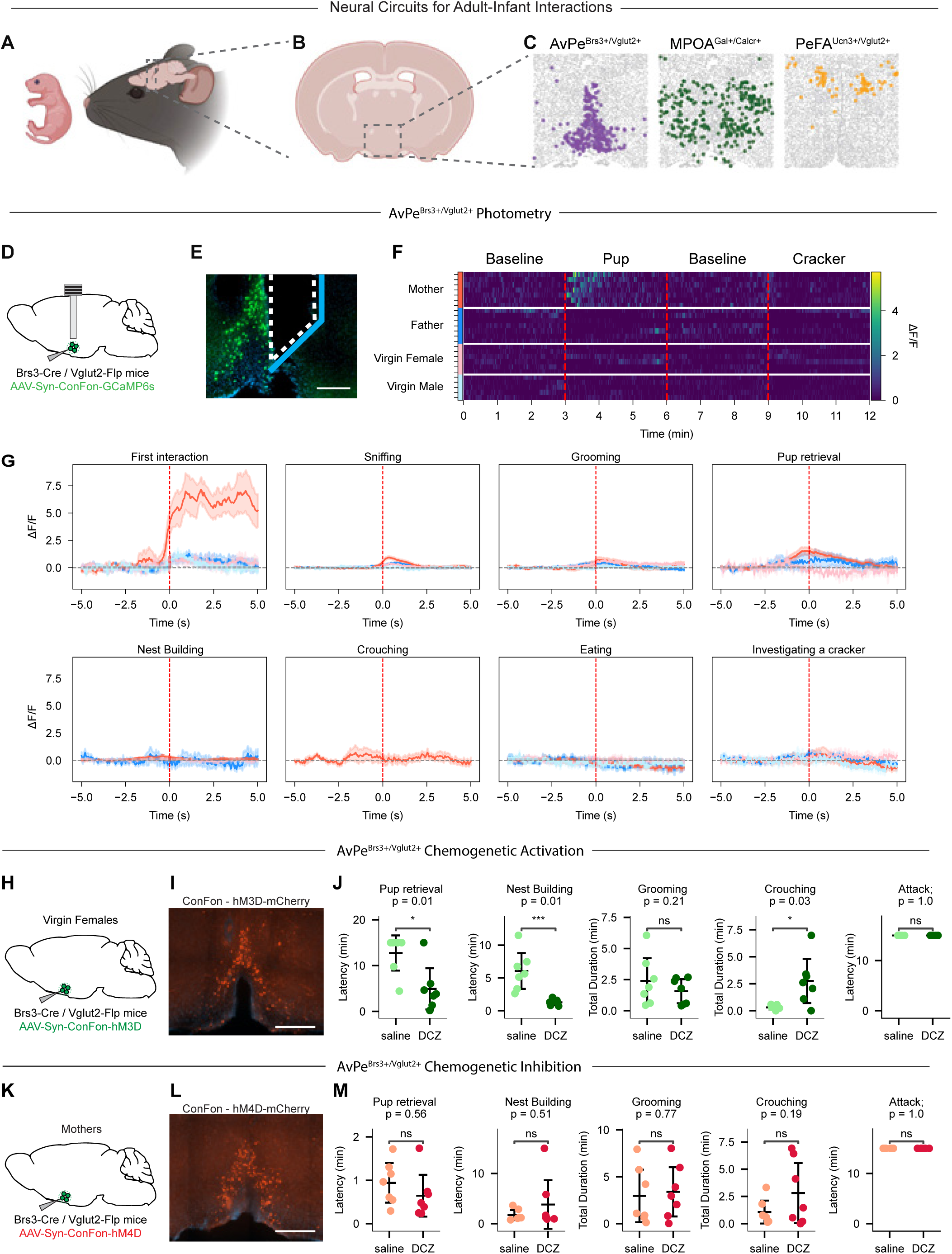
AvPe^Brs3+Vglut2+^ neurons are activate during parenting only in mothers and influence parenting behaviors. (**A**) Cartoon of Adult-Infant interactions. (**B**) Coronal slice of mouse brain highlighting the medial preoptic area as the central hub of Adult-Infant interactions. (**C**) Spatial transcriptomic images depicting three key cell types previously associated with Adult-Infant interactions. (**D**) Experimental procedure for AvPe^Brs3+Vglut2+^ activity imaging. (**E**) Optic fiber placement near GCaMP6m expressing cells in the AvPe. Blue line indicates mirror surface. Scale bar = 200um. (**F**) Heatmap of AvPe^Brs3+Vglut2+^ bulk activity at baseline and during interaction with a pup or a cracker. (**G**) Averaged activity imaging traces from AvPe^Brs3+Vglut2+^ bulk activity during first interaction, sniffing, grooming, pup retrieval, nest building, crouching, eating, and investigating a cracker. Red, mother; pink, virgin female; dark blue, father; light blue, virgin male. (**H**) Experimental procedure for AvPe^Brs3+Vglut2+^ activation in virgin females. (**I**) Expression of chemogenetic activating receptor hM3Dq (red) in AvPe^Brs3+Vglut2+^ neurons. Scale bar = 500um. (**J**) Quantification of behavioral changes following DCZ injection in mice with activated AvPe^Brs3+Vglut2+^ neurons and controls. \**P* < 0.05, \*\*\**P* < 0.001, ns = not significant, Kaplan-Meier survival analysis with log rank test. (**K**) Experimental procedure for AvPe^Brs3+Vglut2+^ inhibition in mothers. (**L**) Expression of chemogenetic inhibiting receptor hM4Di (red) in AvPe^Brs3+Vglut2+^ neurons. Scale bar = 500um. (**M**) Quantification of behavioral changes following DCZ injection in mice with inhibited AvPe^Brs3+Vglut2+^ neurons and controls. ns = not significant, Kaplan-Meier survival analysis with log rank test.

To independently confirm the activity pattern of AvPe^Brs3+Vglut2+^ neurons and further assess their real-time dynamics during behavior, we used an intersectional viral strategy to direct conditional expression of the calcium reporter GCaMP6m in a Cre- and Flp-dependent manner in Brs3-Cre, Vglut2-Flp mutant mice (fig. S1 A,B). Fiber photometry imaging was performed in freely interacting mice (*61*). Because AvPe^Brs3+Vglut2^ neurons lie along the ventricle and are difficult to access, a mirror-tipped fiber was used to achieve maximal optical coverage (Fig. 1 D,E).

AvPe^Brs3+Vglut2+^ neurons displayed a significant increase in activity upon the introduction of pups into the cage of mothers, but not fathers, virgin males nor virgin females (Fig. 1 F,G mothers n=7, fathers n=7, virgin females n=6, virgin males n=5), supporting results obtained with IEGs (*9*). Activity peaked during the first minute of infant interaction and subsequently returned to baseline, revealing minimal activity in pup-related behaviors after the initial interaction. No increase in activity was seen when ad libitum fed animals were eating or investigating a size-matched food item (cheese cracker) (Fig. 1G). Similarly, cumulative activity throughout the duration of the assay was elevated only in mothers during pup presentation, with no changes in other groups during food exploration (fig. S2 A).

The observed pattern of mother-specific pup-induced activity suggests a specific role of AvPe^Brs3+Vglut2+^ neurons in the parenting of mothers. Because parental care by mothers is significantly more robust than in any other parental state, we hypothesized that this excitatory population may enhance the initiation of parental care by boosting the activity of downstream parental circuits common to all parental states. To test this hypothesis, we selectively expressed the excitatory chemogenetic receptor hM3Dq in AvPe^Brs3+Vglut2+^ neurons of virgin female mice (Fig. 1 H,I). In vivo chemogenetic activation of AvPe^Brs3+Vglut2+^ neurons by injecting virgin females with deschloroclozapine (DCZ) (*62*) significantly increased the time spent crouching and decreased the latency to pup retrieval and nest building compared to saline-injected controls (Fig. 1 J n=7 animals). No effect was observed on time spent grooming the pups. Interestingly, although AvPe^Brs3+Vglut2+^ neuronal activity was only detected at the initial phase of parenting, artificial activation of this population strongly increases crouching, a behavior that typically occurs after activity has returned to baseline, suggesting a long-lasting effect of a AvPe^Brs3+Vglut2+^ neuronal activation. Altogether, activation of AvPe^Brs3+Vglut2+^ neurons in virgin females increases several aspects of parental behaviors. By contrast, chemogenetic silencing of AvPe^Brs3+Vglut2+^ neurons in mothers through conditional hM4Di expression and DCZ injection did not notably affect parenting related behaviors, suggesting redundant mechanisms by which parenting behavior in mothers is enhanced (Fig. 1K-M, n=7 animals).

### Accessing cell type-specific programing across parental states

The state- and sex-dependent activity and function of AvPe^Brs3+Vglut2+^ neurons in mothers but not in other parenting states may result from state and sex-dependent differences in connectivity as well as cell intrinsic molecular and biophysical differences. We used single nucleus RNA- and ATAC-seq approaches to uncover how cell type-specific transcriptional programs and chromatin accessibility differ across parenting states and infer specific signaling pathways involved in cellular remodeling across conditions (*63*, *64*). Because the preoptic area is a highly heterogeneous structure containing over 70 molecularly defined neuronal cell types (*9*), we used a Cre-recombinase based approach to enrich for relevant cell populations (Fig. 2A). *Brs3* is expressed in both MPOA^Gal+Calcr+^ and AvPe^Brs3+Vglut2+^ neurons, enabling a Brs3-Cre mouse line to enrich for both populations. To orthogonally access these cell types we also utilized Gal-Cre and Ucn3-Cre lines, chosen to enrich for MPOA^Gal+Calcr+^ and AvPe^Brs3+Vglut2+^ populations respectively. This strategy also led to an enrichment in PeFA^Ucn3+Vglut2+^neurons, a population involved in infanticide, as well as other populations implicated in innate behavioral response.

**Fig. 2.**
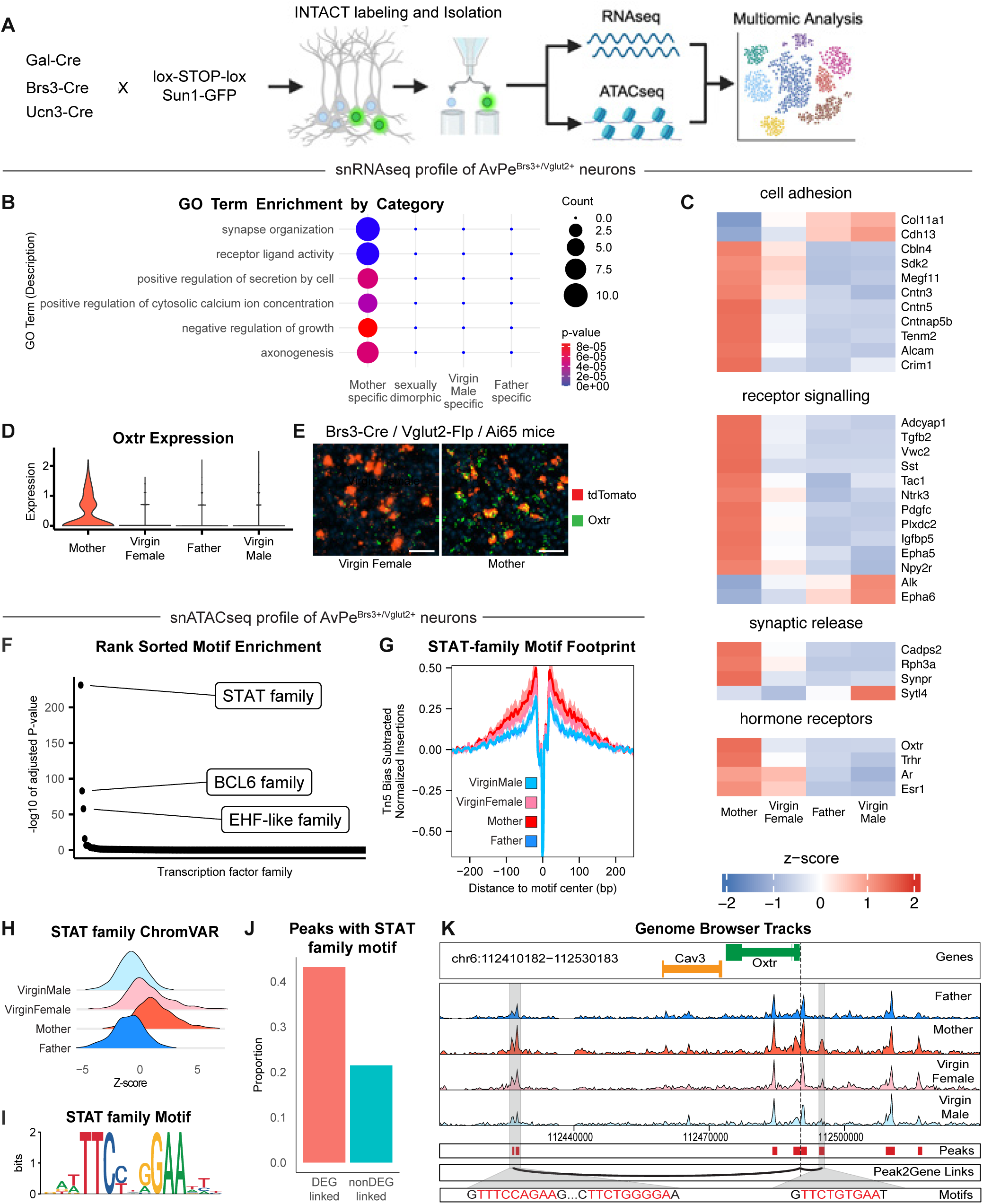
Single cell multi-omic profiling reveals mother-specific transcriptional program influenced by prolactin signaling in AvPe^Brs3+Vglut2+^ neurons. (**A**) Experimental design for enriching dissected POA tissue in specific cell populations and processing multi-omic approaches. (**B**) GO Term enrichment of DEGs in AvPe^Brs3+Vglut2+^ neurons. (**C**) Heatmap of z-scored expression of select DEGs in AvPe^Brs3+Vglut2+^ neurons. (**D**) Oxtr expression across sex and physiological state in AvPe^Brs3+Vglut2+^ neurons. (**E**) Representative image of Oxtr expression (green) and tdTomato expression (red) in AvPe^Brs3+Vglut2+^ neurons. Scale bar = 50um. (**F**) Transcription factor motif families ranked in descending order of enrichment. (**G**) Footprinting score from the STAT-family motif in AvPe^Brs3+Vglut2+^ neurons. (**H**) ChromVAR score of the STAT-family motif. (**I**) Sequence logo of the STAT-family motif. (**J**) Percentage of STAT-family motif containing peaks correlated with gene expression. (**K**) Genome browser plot of ATACseq results at the Oxtr locus in AvPe^Brs3+Vglut2+^ neurons.

Each Cre-driver line was crossed to a Sun1-tagged nuclear reporter strain (*65*) to enable isolation of nuclei tagged in specific cell types (INTACT) from mothers, fathers, virgin females, and virgin males. Samples were split into two pools to be processed separately for RNA and ATAC-seq (Fig. 2A). Rigorous quality control resulted in ∼60,000 nuclei mapped onto 15 previously defined cell types (fig. S3 A,B). Canonical Correlation Analysis (CCA) across RNA-seq and ATACseq modalities shows a high degree of overlap (Fig. S3 C), as does CCA applied across RNA-seq and a previous MERFISH dataset (*9*) (fig. S3 D). This high degree of alignment across modalities allowed for the integration of these three datasets. Specifically, lowly expressed genes detected by MERFISH were imputed onto the RNA-seq dataset, which was then integrated with the ATACseq dataset to allow for multimodal analysis (*66*, *67*).

To identify transcriptional differences across sex and parental state, multiple binary random forest classifiers were trained on the RNA-seq data to determine the information content present across conditions (fig. S3 E) (*68*). Most cell types were invariant across sex and state, including, surprisingly, the infanticide promoting PeFA^Ucn3+Vglut2+^ (*14*, *53*, *60*) which is typically active in virgin males but not in females and mated males, suggesting a cell extrinsic sex- and state-dependent modulation of this cell type activity, presumably through upstream circuit function. By contrast, AvPe^Brs3+Vglut2^ neurons displayed notable differences in all pair-wise comparisons involving mothers (fig S3 E), further instantiating the unique activity profile of this cluster and supporting the notion of cell-intrinsic molecular remodeling of this neuronal population in mothers.

### A mother-unique molecular profile in AvPe^Brs3+Vglut2+^ neurons

To identify transcripts that give rise to mother-specific differences in AvPe^Brs3+Vglut2^ neurons, we performed differential gene expression analysis and hierarchical clustering to evaluate all possible two-way comparisons for this population. This analysis revealed 81 differentially expressed genes (DEGs) across states (fig. S4A). As observed with the classifier, gene expression changes in mothers represented the largest category (75 genes). Characterization of all measured DEGs via gene ontological terminology (GO Term) analysis shows enrichment of specific functional categories exclusively in mothers (Fig. 2B) and differential expression of behaviorally relevant gene families (Fig. 2C) such as cellular/synaptic adhesion members of the cadherin (*Cdh13*) and contactin (*Cntn3*, *Cntn5*, *Cntnap5b*) gene families, suggesting altered connectivity across sex and state. Additionally, several neuropeptides and signaling ligands such as *Tac1*, *Sst*, *Tgfb2*, *Tafa1*, *Pdgfc*, *Ptn*, *Vwc2*, and *Adcyap1* as well as receptors *Ntrk3*, *Epha5*, *Epha6*, *Alk*, *Plxdc2*, and *Npy2r* are differentially expressed along with multiple genes involved in synaptic vesicle release, suggesting differences in neuronal signaling. Finally, the hormone receptors *Oxtr*, *Trhr*, and to a lower extent, *Ar* and *Esr1* were upregulated in mothers, uncovering candidate hormone signals by which sex and state may affect AvPe^Brs3+Vglut2^ neuron function.

Strikingly, transcriptomics data showed that the oxytocin receptor, *Oxtr*, which is a key modulator of parenting behavior (*69–71*), is only expressed in AvPe^Brs3+Vglut2+^ neurons of mothers but not virgin females, fathers or virgin males (Fig. 2D). To confirm these results, Brs3-Cre / Vglut2-Flp mice were crossed to Ai65 reporter mice to yield conditional expression of tdTomato in AvPe^Brs3+Vglut2+^ neurons. AvPe sections from mothers and virgin females were subjected to in situ hybridization to detect transcripts of tdTomato and Oxtr (Fig. 2E). Expression of Oxtr was almost undetectable in virgin females but present in both tdTomato-positive and - negative cells in mothers, confirming the strict mother state-specific expression identified by snRNA-seq analysis.

### Regulation of mother-specific transcription in AvPe^Brs3+Vglut2+^ neurons

We next used chromatin accessibility measurements to interrogate the nature of transcriptional regulation that drives mother specific gene expression in AvPe^Brs3+Vglut2+^ neurons. Although we identified widespread differentially accessible regions (DARs) across sex (fig. S4B), these changes did not correspond to differences in gene expression, suggesting that sexually dimorphic chromatin landscapes do not translate into transcriptional differences in this population at least in adult animals. We next performed transcription factor motif family enrichment analysis to identify candidate signaling pathways underlying the observed changes in gene expression (*72*). Maximum enrichment across all possible pair-wise comparisons revealed remarkably few significantly enriched motifs (Fig. 2F), with the STAT family motif, involved in hormonal signaling, showing the strongest enrichment (enriched >2.8 fold compared to the next most enriched motif). To further investigate the STAT motif across states we performed bias-adjusted footprinting analysis, revealing that chromatin accessibility around STAT motifs is highest in mothers (Fig. 2G). In a similar manner, we used ChromVAR to calculate the deviation in chromatin accessibility scores surrounding the STAT motif compared to expected background accessibility (*73*), also revealing the largest deviation in mothers (Fig. 2H). Although samples were collected from both males and females, and despite the presence of many sex differences in DARs (fig. S4 B), pair-wise comparisons failed to identify any sex differences in motif enrichment but instead revealed the STAT motif as enriched in mother-vs-all comparisons (fig. S4 C).

Stat5b, a member of the STAT family of transcription factors that binds to the canonical STAT motif (Fig. 2I), has been shown to function as the terminal node in prolactin signaling (*74*). Mothers have high circulating prolactin (*75*), which induces the dimerization of the prolactin receptor (Prlr) and facilitates JAK2-mediated phosphorylation of multiple substrates including several kinases that leads to rapid changes in membrane excitability (*74*, *76*). Additionally, Prlr activation induces phosphorylation of the transcription factor Stat5b which leads to long-lasting changes in transcription through the binding to STAT-specific motifs. By analyzing the co-occurrence of gene expression and chromatin accessibility in individual cells from the imputed dataset, it is possible to quantitatively link motif-containing candidate cis-regulatory elements (cCREs) to the genes which they regulate (*67*). Strikingly, DEGs from all possible pair-wise comparisons were linked to cCREs containing one or more Stat5b motifs at twice the percentage of non-DEGs (Fig. 2J). Specific examination of the genomic locus surrounding the Oxtr gene, prompted by the major role played by Oxt signaling in maternal behavior, revealed two cCREs with >0.35 Pearson correlation with Oxtr expression (Fig. 2K). These cCREs each contain canonical Stat5b binding motifs and are more accessible in mothers than any other state, supporting the notion that these elements act as functional enhancers driving increased Oxtr expression in mothers.

### Oxtr signaling in AvPe^Brs3+Vglut2+^ neurons

Our data suggest a model in which elevated circulating prolactin in lactating mothers activates the prolactin receptor in AvPe^Brs3+Vglut2+^ neurons, in turn triggering both immediate membrane effects through ion channels such as Trp-like, L-type, and Ca2+ activated K+ channels (*77–80*) as well as long-lasting transcriptional changes through the Stat5b-mediated pathway in numerous genes including Oxtr (Fig. 3A). To directly test the hypothesis that the sensitivity of AvPe^Brs3+Vglut2+^ neurons to Oxt signaling is enhanced in lactating mothers through Oxtr upregulation, we performed bath application of an Oxtr agonist on AvPe^Brs3+Vglut2+^ neuronal activity in an *ex vivo* AvPe acute brain slice preparation. We crossed Brs3-Cre, Vglut2-Flp mice with the Ai195 line to conditionally express GCaMP7s in AvPe^Brs3+Vglut2+^ neurons and measured intracellular calcium levels as a surrogate marker for neuronal excitability (Fig. 3B). In all imaging sessions combined, we observed low spontaneous activity in most AvPe^Brs3+Vglut2+^ cells (virgin females = 6 animals, 329 cells; mothers = 6 animals 291 cells) (Fig. 3C). Application of the Oxtr-specific agonist (Thr4,Gly7)-Oxytocin (TGOT) to slices prepared from mothers generated dramatic increases in GCaMP7s signal (Fig. 3 C,D), presumably through direct membrane depolarization (*81*). An averaged area under the curve analysis revealed a dramatic increase in activity after TGOT administration in slices prepared from mothers but no change in activity from slices prepared from virgin females, consistent with the absence of *Oxtr* expression in virgin females (Fig. 3E). These results support a model in which mother-specific expression of *Oxtr* in AvPe^Brs3+Vglut2+^ neurons confers sensitivity to oxytocin release in mothers, leading to the activation of this cell type (*82*).

**Fig. 3.**
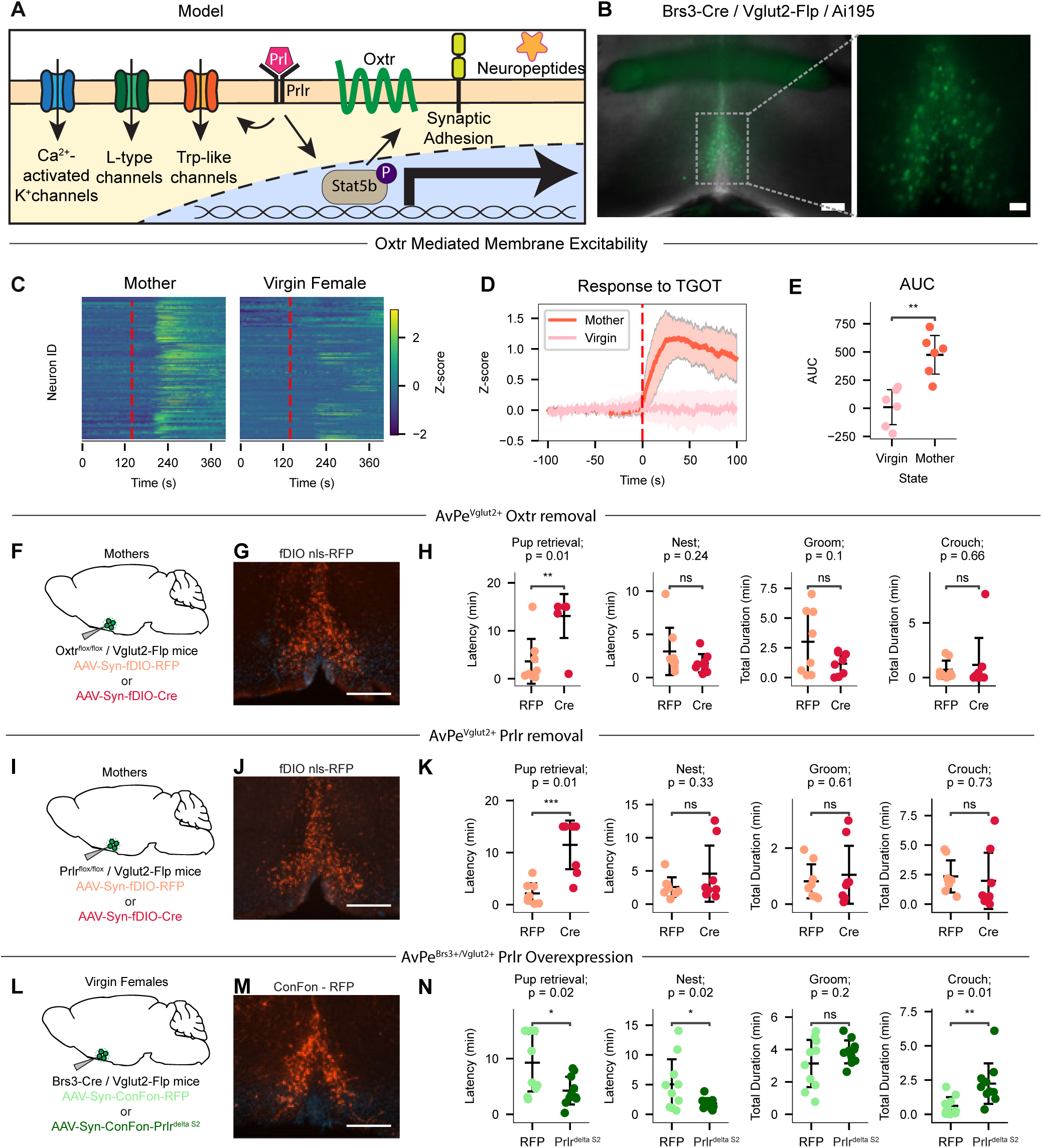
Mother-specific hormone signaling influences AvPe^Brs3+Vglut2+^ neuronal activity and parenting behavior. (**A**) Proposed model depicting prolactin (Prl) activating Prlr to induce both membrane and nuclear effects in AvPe^Brs3+Vglut2+^ neurons, with the later driving cell type-specific activation of Oxtr expression. (**B**) Representative image of GCaMP7s in AvPe^Brs3+Vglut2+^ neurons. Scale bar, zoomed out image = 200um, zoomed in image = 50um. (**C**) Heatmap of AvPe^Brs3+Vglut2+^ neuron activity before and after administration of TGOT. Red line indicates timepoint at which TGOT was added. (**D**) Averaged Z-score of neuronal activity across samples. (**E**) Area under the curve analysis after addition of TGOT. \*\**P* < 0.01, two-tailed t-test. (**F**) Experimental procedure for AvPe^Vglut2+^ removal of Oxtr expression. (**G**) Expression of nls-RFP (red) in AvPe^Vglut2+^ neurons. Scale bar = 500um. (**H**) Quantification of behavioral changes following injection and expression of either Cre-recombinase or RFP control in AvPe^Vglut2+^ neurons. \*\**P* < 0.01, Kaplan-Meier survival analysis with log rank test. (**I**) Experimental strategy for AvPe^Vglut2+^ removal of Prlr expression. (**J**) Expression of nls-RFP (red) in AvPe^Vglut2+^ neurons. Scale bar = 500um. (**K**) Quantification of behavioral changes following injection and expression of either Cre-recombinase or RFP control in AvPe^Vglut2+^ neurons. \*\*\**P* < 0.001, Kaplan-Meier survival analysis with log rank test. (**L**) Experimental strategy for AvPe^Brs3+Vglut2+^ expression of RFP or the constitutively active Prlr^deltaS2^. (**M**) Expression of RFP (red) in AvPe^Brs3+Vglut2+^ neurons. Scale bar = 500um. (**N**) Quantification of behavioral changes following injection and expression of either Prlr^deltaS2^ or RFP control in AvPe^Brs3+Vglut2+^ neurons. \**P* < 0.05, \*\**P* < 0.01, Kaplan-Meier survival analysis with log rank test.

We next sought to investigate if the mother-specific oxytocin signaling in AvPe^Brs3+Vglut2+^ neurons causally contributes to the increased parenting behavior observed in mothers. Due to the lack of an available Cre- and Flp-recombinase conditional Oxtr allele, we used an alternative approach to genetically ablate *Oxtr* expression in excitatory subpopulations of AVPe neurons of adult mice. Mice carrying a conditional Oxtr allele with loxP sites flanking exons 2-3 (*83*) were crossed with a Vglut2-Flp line. Adult females generated by this cross were stereotactically injected in the AvPe with an AAV expressing either Flp-dependent RFP (n = 8), or Flp-dependent Cre recombinase (n = 8) (Figure 3 F,G), leading to the conditional deletion of Oxtr in AvPe^Vglut2+^ neurons. Injected females were mated and tested for parenting behavior ∼3-5 days post-parturition. Compared to RFP controls, Cre-injected animals showed a significantly increased latency to retrieve pups (p = 0.01), while other measures of parental care including nest building, grooming, and crouching were unaffected (Fig. 3H), confirming the role of Oxtr expression in AvPe^Vglut2+^ neurons for normal maternal behavior in postpartum females. Of the four *Vglut2* expressing cell types in the AvPe (fig. S1B), AvPe^Brs3+Vglut2+^ neurons are the only population expressing *Oxtr* or having been described to be involved in parenting (*9*), suggesting that the observed effects are likely mediated through this population, although, we cannot entirely rule out the possibility that manipulating *Oxtr* expression in other AvPe excitatory cell types may also affect parenting behavior.

### Prolactin signaling in AvPe^Brs3+Vglut2+^ neurons

Next, we examined the functional contribution of prolactin signaling in AvPe^Brs3+Vglut2+^ neurons for parenting control. Using a similar approach as described for Oxtr, we crossed Vglut2-Flp animals with mice in which exon 5 of Prlr is flanked with loxP sites (*84*). Adult females were injected with an AAV conditionally expressing Flp-dependent RFP (n = 8) or Flp-dependent Cre recombinase (n = 8) to remove Prlr in the AvPe (Fig. 3 I,J). Cre-injected mothers showed a significantly increased latency to retrieve pups compared to RFP controls (p = 0.01), while other measures of parental care including nest building, grooming, and crouching were unaffected (Fig. 3K). These results suggest that Prlr expression in AvPe^Vglut2+^ neurons is required for the full display of maternal behavior in postpartum females. As with Oxtr removal, we cannot entirely rule out that any of the other three AvPe^Vglut2+^ cell types may be contributing to the observed defect in parenting behavior.

Finally, we assessed whether Prlr signaling in AvPe^Brs3+Vglut2+^ neurons is sufficient to enhance parenting in virgin females. We used the INTERSECT strategy (*85*) to design a novel AAV expressing a constitutively active Prlr^delta^ ^S2^ (*86*, *87*) in a Cre- and Flp-dependent manner (Fig. 3 L,M). Compared to RFP injected controls (n = 10), virgin females with constitutive Prlr signaling in AvPe^Brs3+Vglut2+^ neurons (n = 10) showed a decreased latency to retrieve pups (p = 0.02) and initiate nest building (p = 0.02), as well as an increased total time spent crouching with infants (p = 0.01) (Fig. 3N). These results support our model in which the increased levels of prolactin present in lactating mothers activates Prlr signalling in AvPe^Brs3+Vglut2+^ neurons, leading to expression of Oxtr and enhanced oxytocin sensitivity. This molecular cascade in turn increases neural activity in response to oxytocin release during pup encounters (*82*) and ultimately drives the enhanced parental behaviors characteristic of mothers.

### Core parenting circuits downstream of AvPe^Brs3+Vglut2+^ neurons

The observation that artificial activation of AvPe^Brs3+Vglut2+^ neurons increases parenting behavior in virgin females (Fig. 1J) led us to hypothesize that these neurons may reside upstream of other neuronal populations involved in the control of parenting behavior, such as MPOA^Gal+Calcr+^ neurons (*9*). To assess whether artificial activation of AvPe^Brs3+Vglut2+^ neurons may elicit MPOA^Gal+Calcr+^ activity in the absence of pups, we injected a viral vector to selectively express the chemogenetic activator hM3Dq in AvPe^Brs3+Vglut2+^ neurons in both mothers (n = 8) and virgin females (n = 8) as described earlier (Fig. 1H). Three weeks after viral injection animals were administered either DCZ or saline, and brains were harvested 45 minutes after injection (Fig. 4A). In situ mRNA hybridization was then performed on MPOA sections to assess coexpression of the MPOA^Gal+Calcr+^ marker genes *Gal* and *Calcr* (fig. S5) and the IEG *Fos* as a readout of neuronal activity. Both mothers and virgin females showed a significant increase in *Fos* expression in MPOA^Gal+Calcr+^ neurons following activation of AvPe^Brs3+Vglut2+^ neurons compared to saline injected controls (mother, saline-treated n=235 cells, 4 animals; mothers, DCZ-treated n=348 cells, 4 animals; virgin females, saline-treated n=209, 4 animals; virgin females DCZ-treated n=285 cells, 4 animals; mothers = 7.7% saline, 48.1% DCZ; virgin females = 12.8% saline, 45.6% DCZ; Fig. 4B,C), suggesting that AvPe^Brs3+Vglut2+^ neurons function upstream of this core parenting hub.

**Fig. 4.**
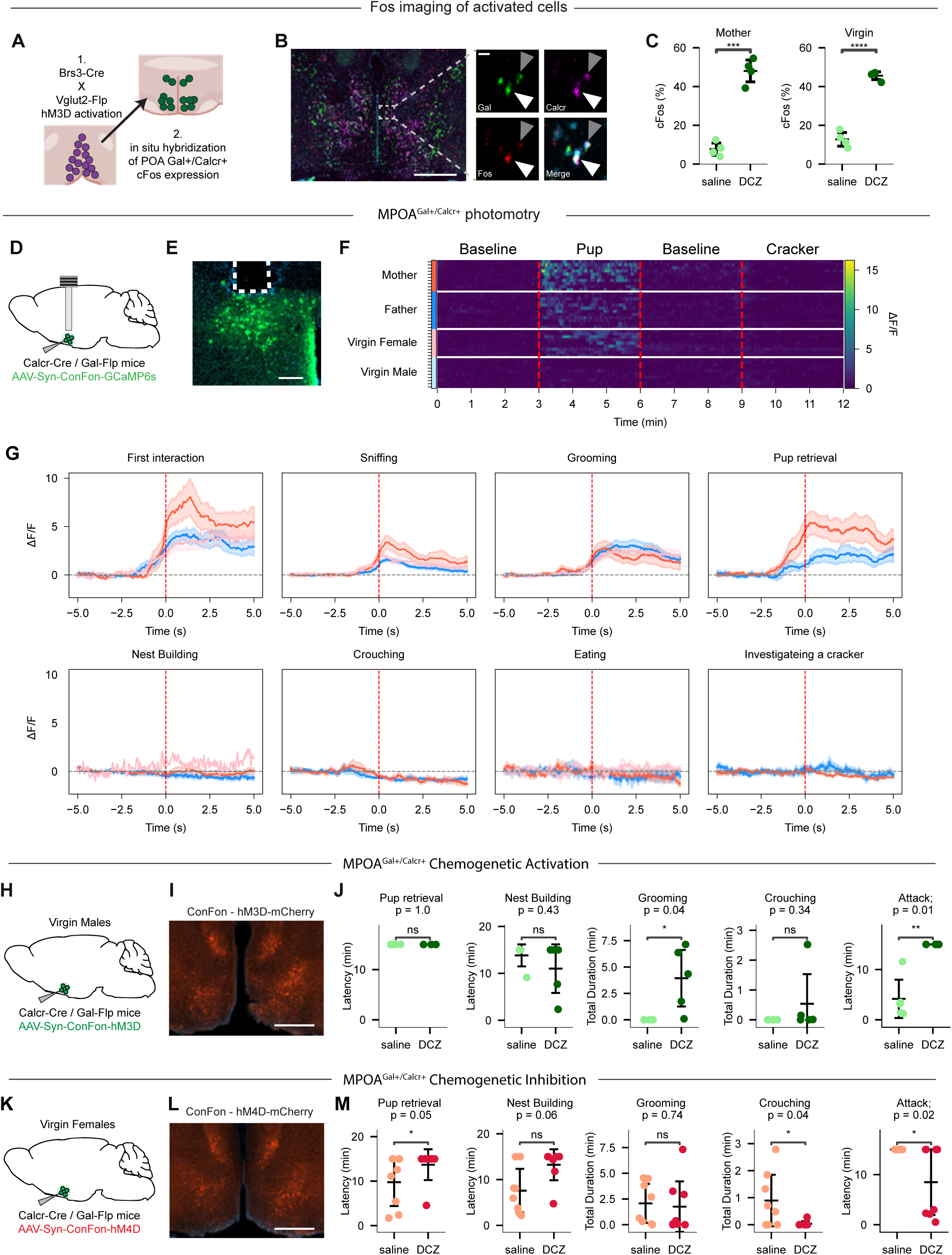
MPOA^Gal+Calcr+^ neurons constitute a core parenting node that acts downstream of AvPe^Brs3+Vglut2+^ neurons. (**A**) Experimental design of AvPe^Brs3+Vglut2+^ neuron activation followed by in situ profiling of MPOA^Gal+Calcr+^ neuron Fos expression. (**B**) In situ expression profiling of Gal (green), Calcr (magenta), and Fos (red) highlighting activated (white arrow) or not activated (grey arrow) MPOA^Gal+Calcr+^ neurons. Scale bar, zoomed out image = 500um, zoomed in image = 10um. (**C**) Quantification of Fos expression following DCZ injection or controls. \*\*\**P* < 0.001, \*\*\*\**P* < 0.0001, two-tailed t-test.(**D**) Experimental strategy for MPOA^Gal+Calcr+^ activity imaging. (**E**) Fiber placement above GCaMP6m expressing cells in the MPOA. Scale bar = 200um. (**F**) Heatmap of MPOA^Gal+Calcr+^ bulk activity during interaction with a pup or a cracker. (**G**) Averaged activity imaging traces from MPOA^Gal+Calcr+^ bulk imaging during first interaction, sniffing, grooming, pup retrieval, nest building, crouching, eating, and investigating a cracker. (**H**) Experimental strategy for MPOA^Gal+Calcr+^ activation. (**I**) Expression of chemogenetic activating receptor hM3Dq (red) in MPOA^Gal+Calcr+^ neurons. Scale bar = 500um. (**J**) Quantification of behavioral changes following DCZ injection in mice with activated MPOA^Gal+Calcr+^ neurons and controls. \**P* < 0.05, \*\**P* < 0.01, Kaplan-Meier survival analysis with log rank test. (**K**) Experimental strategy for MPOA^Gal+Calcr+^ inhibition. (**L**) Expression of chemogenetic inhibiting receptor hM4Di (red) in MPOA^Gal+Calcr+^ neurons. Scale bar = 500um. (**M**) Quantification of behavioral changes following DCZ injection in mice with inhibited MPOA^Gal+Calcr+^ neurons and controls. \**P* < 0.05, Kaplan-Meier survival analysis with log rank test.

Previous studies have shown both real time activity and induction of Fos expression in the broader population of MPOA^Gal+^ neurons during parental care, as well as Fos expression in the MPOA^Gal+Calcr+^ subset more directly involved in parenting (*7–9*). To determine which aspects of parental care are associated with MPOA^Gal+Calcr+^ neural activity, we injected a Cre- and Flp-dependent GCaMP6m expressing AAV into the MPOA of Calcr-Cre, Gal-Flp mice and implanted an optic fiber to monitor neural activity (Fig. 4 D, E). Broad activity was observed in MPOA^Gal+Calcr+^ neurons in the presence of an infant in parental mothers, fathers, and virgin females, but not in non-parental virgin males (Fig. 4F mothers n=10, fathers n=12, virgin females n=9, virgin males n=10). Moreover, by contrast to AvPe^Brs3+Vglut2+^ neurons, which are active during the first moments of pup encounter, MPOA^Gal+Calcr+^ neurons displayed high activity during all pup-directed parenting episodes, but not during non-pup-directed (nest building), passive (crouching) parenting episodes or non-parental behaviors such as investigating or eating a cracker (Fig. 4G).

To further confirm a direct role of MPOA^Gal+Calcr+^ neurons in parenting we sought to determine if functional manipulations would be sufficient to induce parenting behaviors in a cohort of non-parenting virgin males (Fig. 4 H, I). *In vivo* chemogenetic activation of MPOA^Gal+Calcr+^ neurons in virgin females through selective expression of the excitatory chemogenetic receptor hM3Dq led to a significant increase in time spent grooming as well as increased latency to attack in treated animals, while it did not affect retrieval, nest building, or crouching (Fig. 4J n=5 animals), similar to previous reports of optogenetic stimulation of MPOA^Gal+^ neurons in virgin males (*8*). To further assess whether MPOA^Gal+Calcr+^ neurons are required for parental care, we conditionally expressed the inhibitory chemogenetic receptor hM4Di in virgin females (Fig. 4K, L). Strikingly, *in vivo* chemogenetic inhibition of MPOA^Gal+Calcr+^ neurons attenuated pup retrieval and resulted in attack of infants in the majority of animals along with decreased total time spent crouching with pups during the 15 minute assay (Fig. 4M n= 7 animals). These results recapitulate the observed switch from parental to infanticidal behaviors following MPOA^Gal+^ neuron ablation in virgin females (*8*), confirming the specific role of MPOA^Gal+Calcr+^ neurons in parenting behavior.

### Molecular remodeling of MPOA^Gal+Calcr+^ neurons

In view of the role of MPOA^Gal+Calcr+^ neurons in controlling parenting behavior in both males and females, we sought to identify transcriptional differences in the snRNA-seq profiles across parental states, and in particular between parental fathers and non-parental virgin males. Surprisingly, random forest classifier analysis pointed to strong sex differences as well as mother specific transcriptional programs in MPOA^Gal+Calcr+^ neurons but showed a notable lack of differences between parental fathers and non-parental virgin males (fig. S3E). Differential gene expression analysis and hierarchal clustering revealed 260 DEGs, with differences in all possible pair-wise comparisons (fig. S6A,B,C). Despite this complexity, a pattern of sex-dependent expression levels emerged as the largest category (168 genes), followed by genes altered specifically in mothers compared to all other states (56 genes). GO Term enrichment analysis identified gene categories enriched in transcripts showing sex differences as well as genes with mother specific expression patterns (Fig. 5A). Particularly intriguing were the number of terms associated with neuronal functional properties (Fig. 5B). For instance, multiple genes involved in synaptic functions, such as ionotropic neurotransmitter receptors (*Grin2a*, *Grik2*, *Gabrg1*, *Glra1*, *Glra2*, and *Glra3*) and cell adhesion molecules (*Cntn5*, *Cntnap1*, *Clstn2*, *Cdh10*, *Cdh7*, and *Sdk1*), associated with synaptic transmission. Furthermore, differences in neuropeptides (*Nts* and *Gal*) and neuropeptide receptors (*Ntsr1*, *Npy1r*, *Avpr1a*, *Cckbr*, and *Hcrtr1*) suggest that this cluster is subject to differential modulation according to the sex and physiological state of the animal. Additionally, several ion channels including voltage-gated potassium channels (*Kcnab1*, *Kcnc2*, *Kcnh5*, *Kcnh7*, and *Kcnh8*), voltage gated calcium channels (*Cacng3* and *Cacna2d3*), and specialized channels (*Kcnt2*, *Trpm3*, *Asic2*, and *Clca3a1*) were represented in DEGs associated with both sex-differences as well as mother specific expression patterns. Overall, these results highlight a dramatic, and unexpected, repertoire of sex differences and mother specific changes in gene expression in MPOA^Gal+Calcr+^ neurons rather than differences aligning with the differential display of parental care (i.e., mothers, virgin females and fathers vs virgin males).

**Fig. 5.**
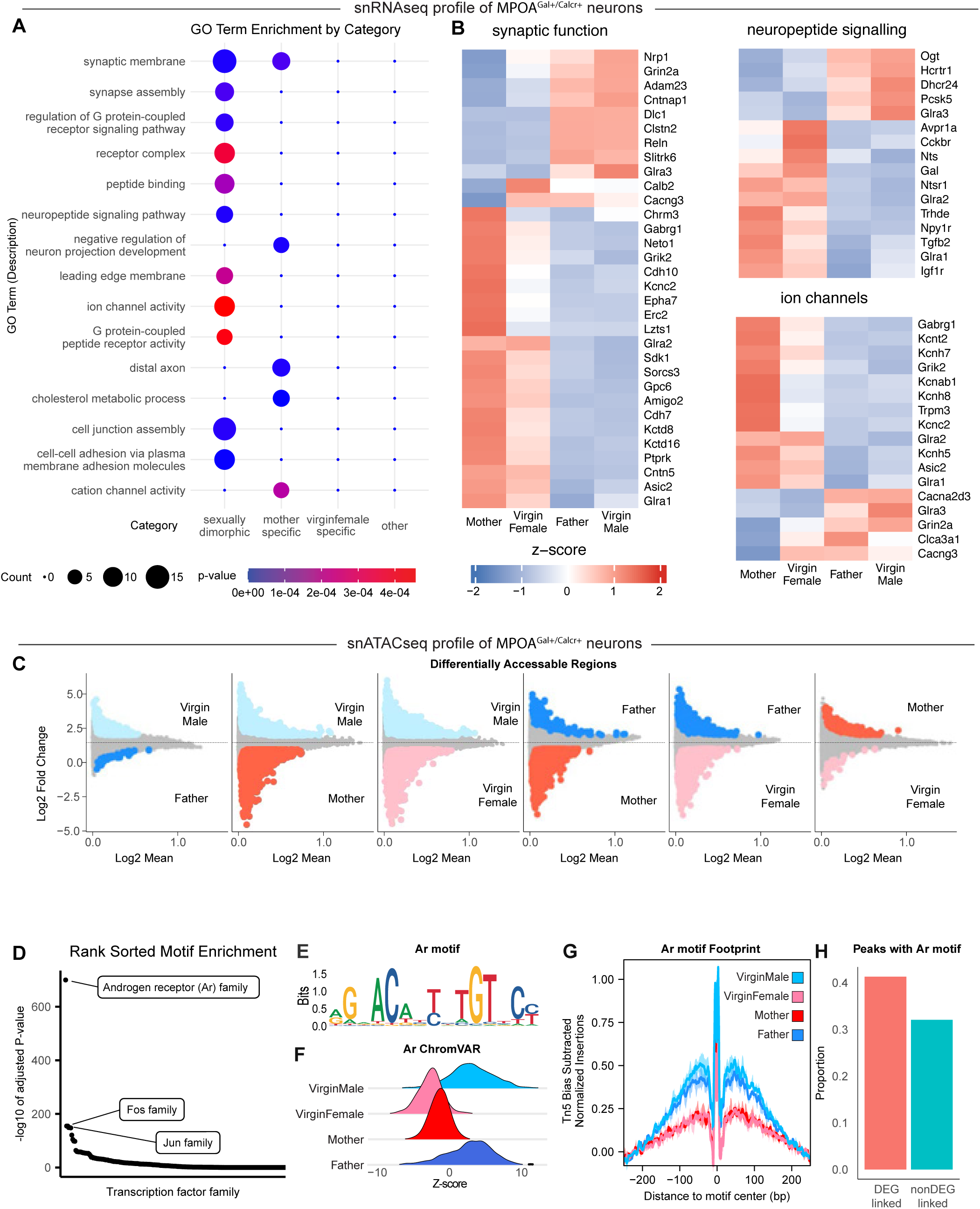
Sex differences in MPOA^Gal+Calcr+^ neurons reveal an androgen based transcriptional program. (**A**) GO Term enrichment of DEGs in MPOA^Gal+Calcr+^ neurons. (**B**) Heatmap of z-scored expression of select DEGs in MPOA^Gal+Calcr+^ neurons. (**C**) Pair-wise log2Fold change plots of differentially accessible regions for all combinations of samples. Color represents significant differences in accessibility. (**D**) Transcription factor motif families ranked in descending order of enrichment in MPOA^Gal+Calcr+^ neurons. (**E**) Sequence logo of the Ar-family motif in MPOA^Gal+Calcr+^ neurons. (**F**) ChromVAR score of the Ar-family motif in MPOA^Gal+Calcr+^ neurons. (**G**) Footprinting score from the Ar-family motif in MPOA^Gal+Calcr+^ neurons. (**H**) Percentage of Ar-family motif containing peaks correlated with gene expression in MPOA^Gal+Calcr+^ neurons.

### Regulation of sex-specific transcription in MPOA^Gal+Calcr+^ neurons

To uncover the specific transcriptional pathways that mediate the observed gene expression programs in MPOA^Gal+Calcr+^ neurons, we examined DARs across sex and parental state. Similar to DEG analysis, the largest number of DARs was observed when comparing animals across sex (Fig. 5C). Examination of transcription factor binding motifs present in accessible regions revealed a dramatic enrichment of the androgen receptor family motif in males (Fig. 5D). This motif is bound by members of the nuclear receptor subfamily 3 group C members: androgen receptor (Ar), glucocorticoid receptor (Gr), progesterone receptor (Pr), and mineralocorticoid receptor (Mr) (*88*). Because levels of both glucocorticoids (*89*) and progesterone (*90*) are higher in females, the opposite of the activity observed here, it is unlikely that these effects are driven by either Gr or Pr. Thus, we may infer that androgens, present at >100x levels in males compared to females (*90*), are driving the enrichment of this motif in males through increased activation of Ar. Moreover, despite numerous transcripts and DARs found to be differentially regulated in either mothers or virgin females, our analysis failed to identify any transcription factor families enriched in their regulation (fig. S7A). As a nuclear hormone receptor, the inactive Ar is sequestered in the cytoplasm by HSP-family chaperones and translocates to the nucleus when bound to androgens such as testosterone or dihydrotestosterone, which are typically higher in males. Once inside nucleus, Ar binds to a conserved DNA motif and influences gene expression (*91*) (Fig. 5E). Intriguingly, the canonical mechanism mediating male-specific sex-differences in the brain involves the conversion of testosterone into estrogen via aromatase, with Esr1 then mediating male-specific expression (*92*). However, by comparison to the dramatic enrichment of Ar binding motifs we see very little enrichment of Esr1 motifs, suggesting that the observed sex differences in gene expression may be mediated directly through Ar.

To measure the ability of Ar to influence chromatin accessibility we calculated both ChromVAR and TF footprinting scores for this motif (Fig. 5F, G). Both metrics reveal that the presence of an Ar family motif is associated with increased chromatin accessibility in males compared to females. To evaluate the contribution of Ar in regulating the observable DEGs, we used our integrated snRNA-seq and snATAC-seq dataset to determine which genomic regions correlate with RNA expression. We then assigned open chromatin regions as cCREs for specific genes and measured the percentage of DEGs linked to cCREs containing Ar family motifs. This comparison revealed that DEGs were more likely to be influenced by cCREs containing Ar family motifs than were non-DEGs (Fig. 5H). These results suggest that androgen signaling through Ar in MPOA^Gal+Calcr+^ neurons influences gene expression in a sex-dependent manner.

### Ar signaling in MPOA^Gal+Calcr+^ neurons

The observed increase in Ar binding in MPOA^Gal+Calcr+^ neurons of males supports a model in which high levels of testosterone activate Ar to influence the expression of many key genes affecting neuronal activity, such as ion channels, synaptic molecules, neuropeptides, and their receptors (Fig. 6A). However, altered combinations of genes often have nonlinear and unpredictable effects on cellular physiology. To test our hypothesis and focus on the cell-autonomous properties related to gene-expression changes in MPOA^Gal+Calcr+^ neurons, we used *ex vivo* slice electrophysiology to assess differences in intrinsic biophysical properties between animals according to sex and state. To exclusively access MPOA^Gal+Calcr+^ neurons, we conditionally expressed mScarlet in Calcr-Cre, Gal-Flp double knock-in animals and performed whole-cell patch clamp of labeled cells in live acute brain slices from mothers, fathers, virgin females, and virgin males (Fig. 6B). Action potential kinetics and inward currents activated by hyperpolarization (Ih currents) in MPOA^Gal+Calcr+^ neurons were unaltered by sex and state (fig. S8A-H). Intriguingly, while resting membrane potential was similar (fig. S8I), we observed a significant increase in membrane resistance in males compared to females (Fig. 6C), mirroring the observed decrease in ion channel expression in males (Fig. 5B). The increase in membrane resistance in males was complemented by an increase in rheobase, or the amount of injected current needed to elicit a single action potential, in cells from parental animals (Fig. 6D). These results paradoxically reflect a scheme in which MPOA^Gal+Calcr+^ neurons in virgin males, which have higher membrane resistance and lower rheobase, should be the most excitable cells, yet this is not seen *in vivo* upon pup encounter (Fig. 4F). To resolve this apparent contradiction, we examined how cells responded to a range of increasing current steps ceasing injections when cells entered depolarization block. When we measured the amount of current a neuron could be injected with before it entered depolarization block, we found striking results. In most animals, including virgin males, a majority of MPOA^Gal+Calcr+^ neurons entered depolarization block quickly (Fig. 6E, G-I). However, MPOA^Gal+Calcr+^ neurons from mothers accepted nearly double the amount of current as other states before entering depolarization block (Fig. 6E, J).

**Fig. 6.**
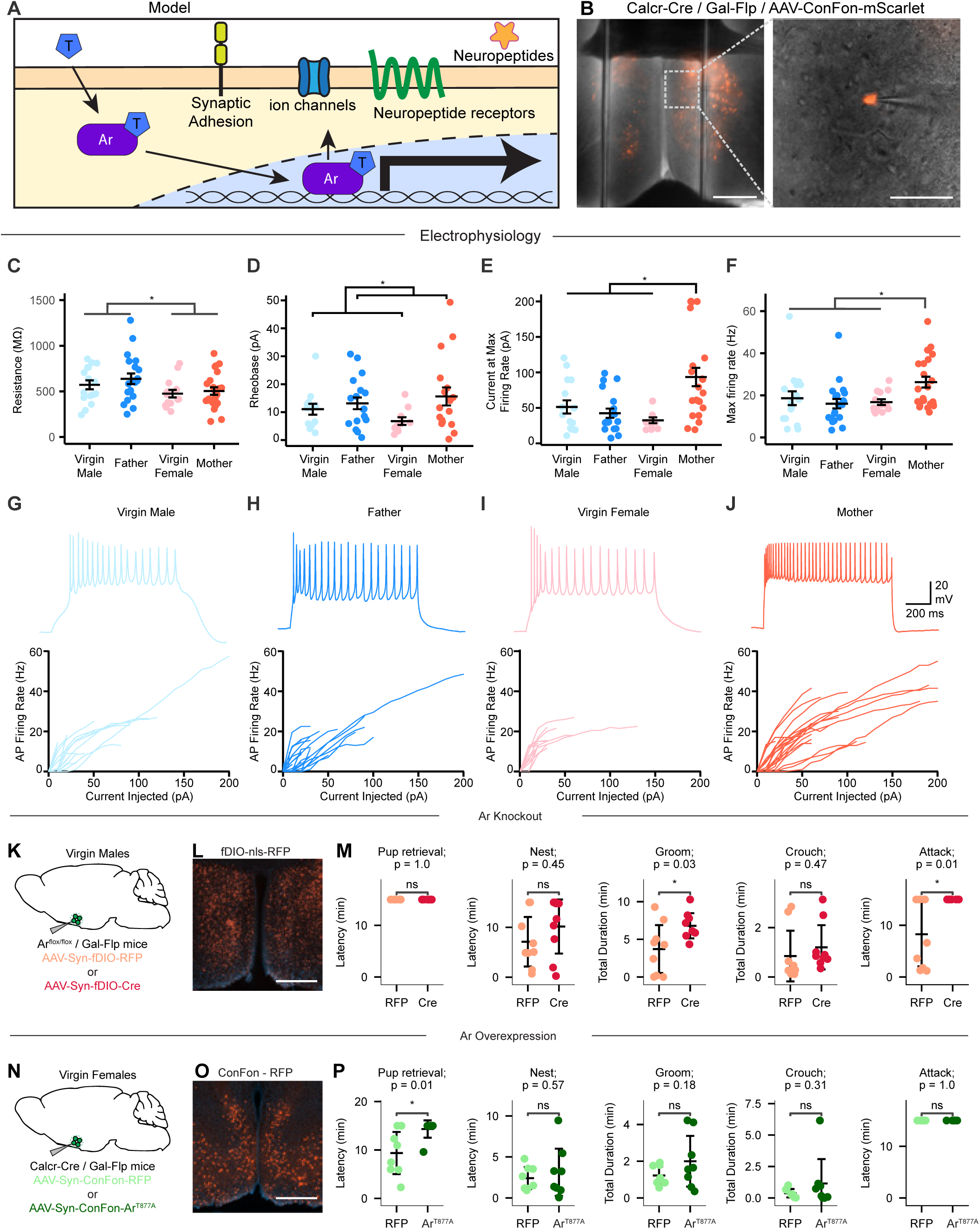
Male-specific hormone signaling influences MPOAGal+Calcr+ neuronal activity and parenting behavior. (**A**) Proposed model depicting activation of the androgen receptor (Ar) by androgen (T) leading to transcriptional changes and differences in ion channel and other gene expression. (**B**) Representative image of MPOAGal+Calcr+ neurons expression mScarlet with patch pipette. Scale bar, zoomed out image = 500um, zoomed in image = 100um. (**C-F**) Representative trace from current clamp experiments from virgin males (**C**), father (**D**), virgin female (**E**), and mothers (**F**) with all samples shown beneath. (**G**) Input resistance of cells. Two-way ANOVA, Sex Effect: F(1,69) = 5.391, p = .0232, virgin male n = 16, father n = 20, virgin female n = 14, mother n = 23. (**H**) Rheobase of cells (Two-way ANOVA, State Effect: F(1, 54) = 4.482 p = .0389, virgin male n = 13, father n = 18, virgin female n = 10, mother n = 17). (**I**) Maximum firing rate (Hz) achieved before depolarization block. Two-way ANOVA, Interaction Effect: F(1,67) = 7.719, p = .0071, Sex Effect: F(1,67) = 4.122, p = .0463; Tukey’s multiple comparisons, mother vs. virgin female p = .0452, mother vs. father p = .0020, mother vs. virgin male p = .1095, virgin male n = 16, father n = 19, virgin female n = 13, mother n = 23.(**J**) Current being injected when a cell reached its maximum firing rate. Two-way ANOVA, Interaction Effect: F(1, 65) = 10.80 p = .0016; State Effect: F(1, 65) = 6.014 p = .0169; Tukey’s multiple comparisons, mother vs. virgin male p = .0215, mother vs. father p = .0016, mother vs. virgin female p = .0009, virgin male n = 15, father n = 19, virgin female n = 12. (**K**) Experimental procedure for MPOAGal+ removal of Ar. (**L**) Expression of nls-RFP in MPOAGal+ neurons. Scale bar = 500um. (**M**) Quantification of behavioral changes following injection and expression of either Cre-recombinase or RFP control in MPOAGal+ neurons. *P < 0.05, Kaplan-Meier survival analysis with log rank test. (**N**) Experimental procedure for MPOAGal+Calcr+ expression of the constitutively active of ArT877A. (**O**) Expression of RFP in MPOAGal+Calcr+ neurons. Scale bar = 500um. (**P**) Quantification of behavioral changes following injection and expression of either ArT877A or RFP control. *P < 0.05, Kaplan-Meier survival analysis with log rank test.

Additionally, the maximal firing rate of these cells was increased compared to other states (Fig. 6F). Together, these results suggest that sex and state synergize to create unique physiological states in MPOA^Gal+Calcr+^ neurons. Decreased input resistance in females, potentially driven by differential Ar signaling, sets the stage for a more dynamic response to input, which may be taken full advantage of in mothers. MPOA^Gal+Calcr+^ neurons in mothers are uniquely dynamic in their response to incoming signaling, in line with additional mother-specific changes in gene expression (Fig 5A), as they allow a broad range of responses over stronger inputs with the ability to give stronger output.

In view of the apparent outsized influence of Ar in mediating both molecular and biophysical differences in MPOA^Gal+Calcr+^ neurons, we next sought to assess the role of Ar in this neuronal population in behaving animals. From the published literature, sex differences in gene expression may impart functional differences between males and females or may serve to compensate for other physiological differences and in fact minimize sex differences (*93*). To assess the role of Ar activity in MPOA^Gal+Calcr+^ neurons of adult mice, and due to the lack of a double conditional allele to perform intersectional gene deletion, we used a conditional Ar allele in which exon 2 is flanked with loxP sites (*94*) crossed with a Gal-Flp animal. This allows for normal development as the Ar gene will remain intact until the expression of Cre-recombinase. To functionally remove Ar in adult animals, adult virgin males were stereotactically injected with an AAV vector mediating Flp-dependent expression of RFP (n = 8) or Cre recombinase (n = 8) into the MPOA (Fig. 6 K, L). Three weeks after surgery, animals were assayed for pup-induced behaviors. AAV-mediated ablation of Ar in adult virgin males resulted in a significant increase of the total time spent grooming an infant while infant attack was abolished, and no substantial differences in latency to build a nest or the total duration spent crouching with the infant were observed (Fig. 6M), suggesting that ablation of Ar signaling in MPOA^Gal+^ neurons of virgin males results in significant increase in parental care and decrease in infant mediated aggression.

To determine if Ar signaling in MPOA^Gal+Calcr+^ neurons is sufficient to decrease parental care, we engineered a novel AAV mediated construct. Drawing on observations from prostate cancer, where Ar mutations produce constitutively active receptors (*95*, *96*), we generated a Cre- and Flp-dependent Ar gene containing the activating T877A mutation. Calcr-Cre, Gal-Flp virgin female mice were injected with a Cre/Flp dependent RFP (n=8) or constitutively active Ar^T877A^ (n=8) and allowed three weeks for recovery and expression (Fig. 6 N, O). Compared to control animals, expression of Ar^T877A^ resulted in a significant reduction in animals that retrieve pups to their nest (Fig. 6P). Together with deletion experiments, these results suggest a model in which high level of androgens activates Ar signaling, leading to a reduction of MPOA^Gal+Calcr+^ neuronal activity and decreased parenting behavior.

## Discussion

Here, we provide a cell type–resolved molecular dissection of hormonally regulated circuits underlying infant-directed behaviors, identifying the specific neuronal populations that serve as targets of hormonal control and characterizing cell-intrinsic transcriptional and biophysical changes that drive state- and sex-specific expression of parental care. These findings provide a new mechanistic framework for hormonal regulation of this essential social behavior with high resolution (Fig. 7). By integrating intersectional genetics, activity imaging, and manipulation with single-cell multi-omics and electrophysiology, we show that two hypothalamic preoptic area cell types, the excitatory AvPe^Brs3+Vglut2+^ population and the inhibitory MPOA^Gal+Calcr+^ population, are molecularly and biophysically tuned in a sex- and state-dependent manner to recalibrate circuit activity and, ultimately, parental behavior. Specifically, we identify a unique prolactin-driven regulatory population in mothers, in which a hormonal cascade cell-autonomously tunes the molecular profile and activity to boost downstream parenting circuits, and we uncover a second, androgen-sensitive population that modulates parenting in a sex- and state-dependent manner. Mothers exhibit rapid activation in AvPe^Brs3+Vglut2+^ neurons when encountering pups, and chemogenetic excitation of these cells in virgin females lowers retrieval latency and increases crouching, whereas silencing in mothers is behaviorally redundant, implying parallel facilitatory pathways in the maternal brain. Single-nucleus RNA- and ATAC-seq further reveal a prolactin-dependent STAT5b transcriptional program that induces *Oxtr* expression specifically in the maternal state. *Ex vivo* calcium imaging confirms that AvPe^Brs3+Vglut2+^ neurons in maternal brains gain oxytocin sensitivity, linking molecular changes to biophysical properties. Intriguingly, although chemogenetic inhibition fails to alter parental care, conditional loss of either *Oxtr* or the prolactin receptor delays retrieval in mothers, suggesting potential circuit changes associated with genetic removal of key hormonal signaling pathways. Supporting the importance of these pathways, constitutive prolactin-receptor signaling in virgins is sufficient to stimulate caregiving.

**Fig. 7.**
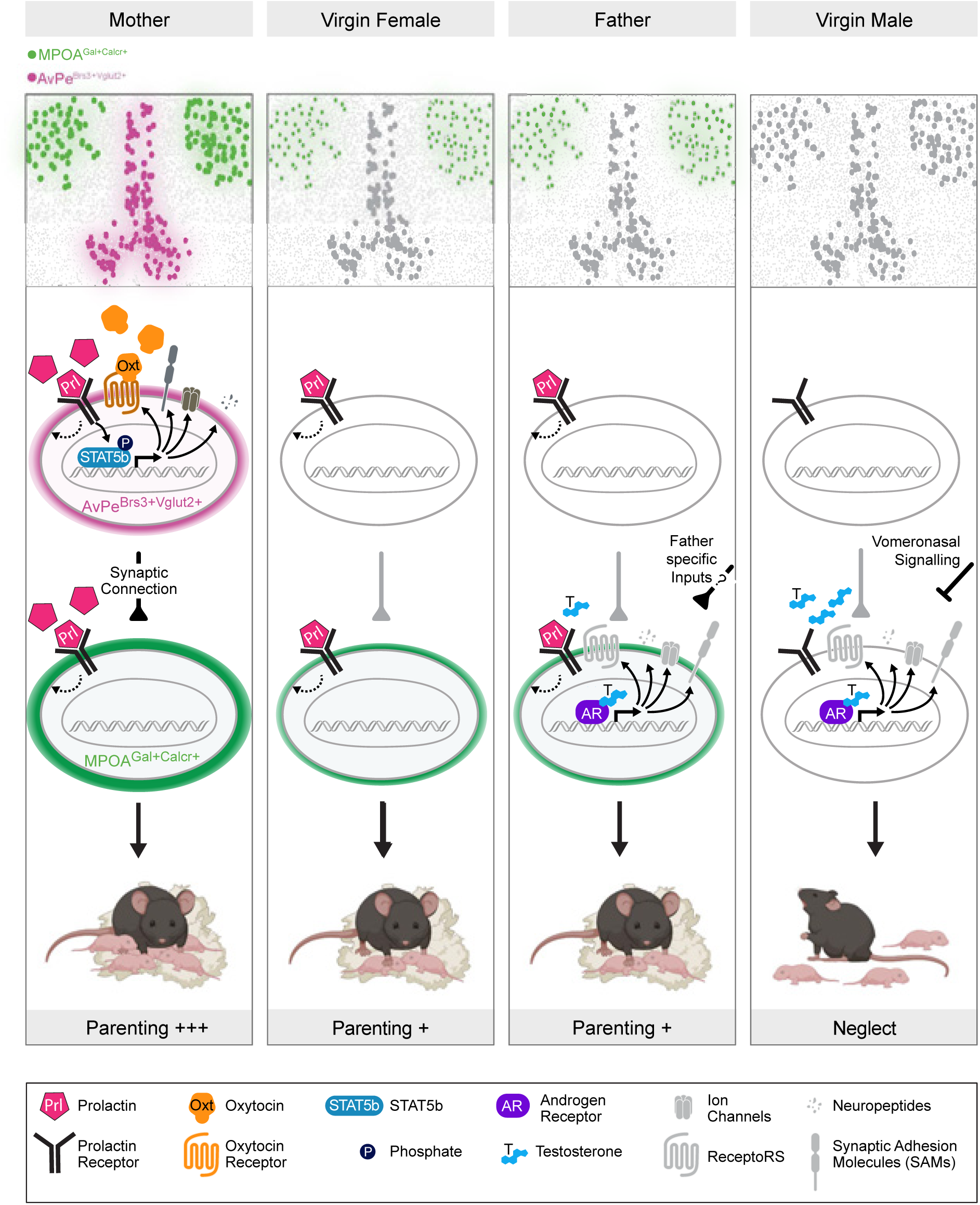
Model of hormonal influence on neural circuits controlling parenting behavior. AvPe^Brs3+Vglut2+^ and MPOA^Gal+Calcr+^ neurons control parenting and are influenced by exposure to circulating hormones. Lactogens such as prolactin bind to Prlr and exert both immediate and long-lasting transcriptional effects that results in mother-specific gene expression patterns of multiple classes of neuronally relevant molecules to increase responsiveness to oxytocin. Androgens such as testosterone activate Ar to alter the expression of several biomolecules including ion channels which inhibit neuronal activity. Fathers potentially overcome this molecular inhibition through father-specific inputs, while virgins males receive inhibitory signaling from VNO circuits.

AvPe^Brs3+Vglut2+^ activation recruits MPOA^Gal+Calcr+^ neurons, which function as a bidirectional hub that orchestrates caregiving versus infanticide. Real-time neural activity monitoring shows MPOA^Gal+Calcr+^ activity scales with pup handling across mothers, fathers, and parental virgins but is muted in infanticidal males, and chemogenetic experiments demonstrate that MPOA^Gal+Calcr+^ neurons are both necessary and sufficient to bias behavior toward parenting or attack. Chromatin accessibility and transcription factor motif analyses identify androgen-receptor signaling as the dominant driver of male-biased differentially accessible regions in these neurons, with Ar-linked enhancers controlling a large subset of ion-channel genes; this transcriptional program is mirrored by higher membrane resistance in males. Removing Ar in MPOA^Gal+^ neurons abolishes pup attack in virgin males, whereas virally introducing a constitutively active Ar mutant into virgin females suppresses retrieval, demonstrating causal necessity and sufficiency. This creates an unexpected dissociation in males: fathers activate MPOA^Gal+Calcr+^ neurons during parenting, yet these neurons show little father-specific transcriptional remodeling relative to virgin males. Instead, a previous study may point toward a circuit regulation lying upstream of MPOA^Gal+Calcr+^ neurons. The vomeronasal organ (VNO), involved in pheromone detection, is strongly activated in virgin males after exposure to pups but remain silent in fathers (*97*). This suggests that changes in circuit activity upstream of MPOA^Gal+Calcr+^ neurons may mediate their activation in fathers, potentially through recruitment of father-specific inputs. Additionally, Prlr has been implicated in father-specific parenting behavior, although with mixed implications (*98*).

Taken together, these data support a model in which hormone-gated transcriptional tuning reconfigures intrinsic excitability and receptor repertoires in discrete nodes of a parenting circuit, thereby biasing information flow according to internal state. Prolactin sensitizes an upstream excitatory trigger in mothers, and androgens dampen a downstream hub in males, jointly determining whether infant cues evoke nurturing or aggressive responses. This mechanism reconciles long-standing observations that parental circuits are latent in both sexes yet expressed according to reproductive context and that maternal behavior can be induced in virgins by hormonal treatments.

Beyond parenting, the logic uncovered here, that systemic hormones targeting cell type-specific enhancers to shape network activity, may generalize to other social and instinctive behaviors such as mating, defense, or feeding, where internal state must dynamically reprioritize behavioral repertoires. Moreover, while we focused on prolactin and androgens, other signals likely converge on the same enhancers, and parsing their combinatorial logic will require multiplexed perturbations.

In sum, by charting how hormonal cues enact cell type-specific transcription to recalibrate neural excitability, we move from descriptive circuit maps toward a mechanistic hormone-to-enhancer-to-behavior framework. This blueprint not only deepens our understanding of parental behavior but also highlights potential entry points for therapeutic modulation of postpartum disorders or pathological aggression.

## Resource availability

### Lead contact

Further information and requests for resources should be directed to the lead contact, Catherine Dulac.

### Data and code availability

https://github.com/Dulac-Lab/Logeman_et_al,

## Acknowledgments

We thank Stacey Sullivan for assistance with animal husbandry and logistics, Samantha Finkbeiner and Sarah Craig for technical assistance, MCB Graphics for help with illustrations, members of the Dulac lab for helpful feedback on our manuscript, the Harvard Center for Biological Imaging (RRID:SCR_018673) for infrastructure and support, and The Bauer Core Facility at Harvard University for assistance with sequencing.

## Author contributions

B.L.L. and C.D. conceived the study; B.L.L. and P.M.H. performed experiments with help from M.T., V.K., and H.S.K; B.L.L. and P.M.H. analyzed data; B.L.L., P.M.H., and C.D. wrote the manuscript, with input from M.T., V.K., and H.S.K.

## Materials and Methods

### Animals

Mice were maintained on a 12 h:12 h dark-light cycle with access to food and water ad libitum. The temperature was maintained at 22 °C and the humidity between 30% and 70%. All experiments were performed in accordance with National Institutes of Health guidelines and approved by the Harvard University Institutional Animal Care and Use Committee. The Gal::Cre BAC transgenic line (Stock: Tg(Gal-cre)KI87Gsat/Mmucd, 031060-UCD) and Ucn3::Cre BAC transgenic line (STOCK Tg(Ucn3-cre) KF43Gsat/Mmcd 032078-UCD) were imported from the Mutant Mouse Regional Resource Center and maintained on a C57BL/6 J background for >10 generations. The INTACT reporter mouse B6.Cg-Brs3tm1.1(cre/GFP)Rpa/J, B6;129-Gt(ROSA)26Sortm5(CAG-Sun1/sfGFP)Nat/J, the Cre- and Flp-dependent GCaMp7s mouse B6;129S6-Igs7tm195(tetO-GCaMP7s,CAG-tTA2)Tasic/J, the Cre- and Flp-dependent tdTomato reporter mouse B6;129S-Gt(ROSA)26Sortm65.1(CAG-tdTomato)Hze/J, and the Oxtr floxed mouse B6.129(SJL)-Oxtrtm1.1Wsy/J were obtained from The Jackson Laboratory. Calcr-IRES-Cre mice were generated by the Gene Targeting and Transgenics Center at the Janelia Research Campus using accession number NM_001042725.1. The androgen receptor floxed mouse was obtained from W. Walker and the prolactin receptor floxed mouse was obtained from R. Banerjee.

### Constitutively active receptor design and production

To create the Prlr^delta^ ^S2^ residues 126-224, corresponding to the S2 domain, of the wild type mouse Prlr (Uniprot Q08501) were omitted from a synthesized gBlock (IDT). Similarly, mouse Ar (Uniprot P19091) was synthesized as a gBlock (IDT) with residue 857 corresponding to alanine rather than threonine to create Ar^T877A^. Both genes were then subjected to the optimized INTERSECT protocol(*85*) to create Cre- and Flp-recombinase dependent constructs in the pAAV vector. Custom Prlr^delta^ ^S2^ and Ar^T877A^ AAVs of the PHP.eB serotype were produced using an optimized vector production protocol(*99*). The average yield was ∼ 1×10^12^ vg per plate.

### Microscopy

#### RNA in situ hybridization

Fluorescence in situ hybridization experiments were performed using the RNAscope Assay V2 kit (Advanced Cell Diagnostics). Brains were harvested and frozen in optimal cutting temperature medium, then sliced on the cryostat at a thickness of 16 μm. The RNAscope Assay was performed according to the manufacturer’s instructions using probes designed against indicated transcripts. Slides were imaged on an Axio Scan.Z7 microscope at ×20. Cells were counted manually by an investigator blinded to the sample names.

##### Immunofluorescent and fluorescent protein imaging

Animals were perfused with PBS followed by 4% PFA after deep anesthesia by isoflurane. Brains were dissected out and placed in 4% PFA for 1–2 nights, then vibratome sectioned at a thickness of 60 μm. For cFos staining, floating coronal sections were washed three times (5 min each time) in 1× PBS with 0.3% Triton X-100 (PBST), then blocked for 1 h in Animal Free Blocker 5× Concentrate (Vector Laboratories) diluted to 1× in PBST. All washes and blocking were performed at room temperature with gentle shaking at 100 rpm. Sections were then placed in blocking solution at 1:1,000 dilution with the primary rabbit anti-Fos (9F6, Cell Signaling) and left overnight with gentle shaking at 4 °C. Sections were again washed three times for 5 min each in PBST, then incubated in blocking solution at 1:10,000 dilution with secondary antibody Goat anti-Rabbit IgG Alexa Fluor™ 568 (A-11011, Invitrogen) for 2 h. Following three more washes as before, individual sections were then mounted onto a microscopy slide with VECTASHIELD (Vector Laboratoires) and kept at room temperature for at least one night before imaging. Sections were imaged at ×10 using an Axio Scan.Z7 microscope. Cells were counted manually by an investigator blinded to the sample names.

### In vivo fiber photometry

#### Viral injection and fiber implantation

All surgeries were performed under aseptic conditions with animals anesthetized with isoflurane (1–2% at 0.5–1.0 l min −1). Analgesia was administered pre-(buprenorphine, 0.1 mg kg−1, intraperitoneal) and post-operatively (ketoprofen, 5 mg kg −1, intraperitoneal). To express the calcium sensor GCaMP6m (Addgene), we injected 500nl of virus solution bilaterally into the medial preoptic hypothalamus. For targeting AvPe^Brs3+Vglut2+^ neurons we used the following coordinates: ML=0.15, AP=0.55, DV=-5.2. For targeting MPOA^Gal+Calcr+^ neurons we used the following coordinates: ML=0.35, AP=-0.1, DV=-4.8. We then implanted an optic fiber (200μm diameter, Doric Lenses) slightly above the virus injection site and affixed to the skull with dental cement. For targeting AvPe^Brs3+Vglut2+^ neurons a mirror-tipped fiber was used, while for targeting MPOA^Gal+Calcr+^ neurons a flat tipped fiber was used. The implanted mice were housed singly for 1 week to recover before being either being co-housed with an animal of the opposite sex to mate or remaining in an isolated cage for an additional 3 weeks.

#### Calcium imaging and behavioral assays

All behavioral experiments were performed during the dark cycle of the animals in a room illuminated by infrared and red light. Before recording, a magnetic patch cord (200 µm diameter, NA=0.37, 3m long, Neurophotometrics) was connected to the optical fiber implanted on the head of the animal in its home cage and allowed for habituation to the new environment for 30 min.

Once the recording started, the patch cord simultaneously delivered excitation light at 415 nm and 470 nm wavelengths (Neurophotometrics, FP3002) and collected fluorescence emissions. Photometry recordings and behavior video acquisition were synchronized using a common TTL input. To begin the experiments, mice were recorded for ∼5 min without any stimuli to obtain a baseline recording. Then two C57BL/6 J pups, 1–3 days of age, were introduced to the animal’s cage in each corner furthest from the nest and the adult was allowed to interact with the infant for ∼15 min. The infants were then removed and ∼5 min later a cheese cracker was introduced into the cage and recorded for ∼5 min. Behavior videos were scored manually using the Boris open-source software by a scorer blinded to animal identity. The following behavioral modules were scored: first interaction with pup, pup sniffing, pup grooming, pup retrieval to the nest, nest building, crouching (animal hovers above the pup in the nest), eating the cracker, and investigating the cracker.

#### Fiber photometry data analysis

Analysis was performed using custom Python scripts built from established workflows previously described(*100*). Only recordings with a stable baseline were included. A lowpass filter was first applied to the raw signal over each entire recording session to reduce noise, followed by photobleaching correction using a double exponential fit before finally correcting for movement artifacts by subtracting a linear fit of the isosbestic channel. Data were then Z-scored for comparison. Individual behaviors within a session were averaged and these averages were compared across animals of all states.

### Chemogenetics

Viral injection was performed in the same manner as for calcium imaging with either a ConFon excitatory hM3Dq (Addgene) or inhibitory hM4Di (Addgene) DREADD. Animals were allowed 3-4 weeks to recover before behavioral assays. On the day of the experiment, animals were intraperitoneally injected with either saline or DCZ (100 μg per kg) 15 min before introduction of two C57BL/6 J pups and recorded for 15 min.

### Single-nucleus sequencing

#### Behavioral screening and harvest

All animals used for sequencing were subjected to behavioral screening to ensure that either pup retrieval or attack was observed. Before behavioral testing animals were housed individually for 5–7 days unless otherwise specified. Experiments started at the beginning of the dark phase and were performed under dim red light. Testing was performed in the home cage and preceded by a 2 hour habituation period with food pellets and hydrogel present. Two 1–4-day-old C57BL/6 J pups were placed in different corners opposite the nest. Once retrieval/attack occurred a timer was started and animals harvested 5 min later for immediate dissection of the anterior hypothalamus. Tissue was then snap frozen in liquid nitrogen and stored at −80 degrees.

#### Tissue processing

Dissected tissues from 5 mice were pooled for each sample, dounce homogenized on ice and centrifuged at 500g for 5 min at 4 °C Supernatant was removed and passed through a 70-μm filter then through a 20-μm filter (MACS Smart strainer) to remove large debris. DAPI was added to the suspension and nuclei were isolated via FANS. Following centrifugation, nuclei were split into separate pools to undergo processing via Chromium Next GEM Single Cell 3ʹ Reagent Kits v3.1 and Chromium Next GEM Single Cell ATAC v2 protocols from 10x Genomics separately. The resulting suspensions were loaded into a 10x Genomics Chromium single-cell chip with the aim of sequencing 7,000–8,000 nuclei per sample. The libraries were sequenced on an Illumina NovaSeq 6000 instrument using instructions provided by 10x Genomics. Paired-end sequencing with read lengths of 150 nt was performed for all samples.

Illumina sequencing reads were aligned to the mouse genome using the 10x Genomics CellRanger pipeline with the default parameters.

#### snRNA-seq analysis

Illumina sequencing reads were aligned to the mouse genome using the 10X Genomics CellRanger ARC pipeline with default parameters. For initial analysis, we relied on the R package Seurat and standard data analysis practices. We filtered out nuclei with more than 400 or fewer than 100,000 UMIs, with fewer than 250 genes and with more than 20% of UMIs belonging to one gene. Although rare, nuclei with more than 10% mitochondrial or ribosomal UMIs and more than 1% IEG, apoptotic or red blood cell UMIs were also filtered out. We defined the main cell classes (glia, neurons) and separated out inhibitory and excitatory neurons using Seurat clustering and known marker gene enrichment.

#### Cell type assignments

Cell types were defined separately among inhibitory and excitatory neurons using the same procedure. For every individual dataset, SCTransform was first used to normalize all cells across sex and physiological state, regressing out percentages of mitochondrial UMIs, ribosomal UMIs and largest gene. After PCA cells were embedded in a K-nearest neighbor graph and subsequently clustered through the Louvain algorithm. Each cell from this dataset was then mapped to the MERFISH reference atlas from Moffit et al, using canonical correlation analysis-based (CCA) label transfer. If the average top score for each cluster was >0.4 the cluster was assigened the cell type from the Moffit et al dataset.

#### snRNA-seq imputation

As not all genes were detected with the snRNA-seq, a select few genes were imputed via alignment with the Moffitt et MERFISH dataset. For the AvPeBrs3+Vglut2+ and MPOAGal+Calcr+ populations, each unique sex and/or state was subset and integrated with the corresponding sex and/or state using the previously mentioned CCA alignment. From there, a k-nearest neighbor graph was constructed to identify MERFISH cells that most closely resembled the snRNA-analyzed cells. For genes which were determined to be differentially expressed in the MERFISH dataset but not the snRNA-seq dataset, the MERFISH gene expression values were then transferred to the snRNA-seq dataset.

#### snATAC-seq analysis

snATAC-seq profiles were filtered to include only nuclei with at least 500 fragments. Cell type-specific peaks were called for AvPeBrs3+Vglut2+ and MPOAGal+Calcr+ populations using Genrich with default parameters. To compile a cell type-specific union peak set, we combined peaks from all sexes and physiological states. Overlapping peaks were then handled using an iterative removal procedure. First, the most significant peak, that is, the peak with the smallest P value, was kept and any peak that directly overlapped with it was removed. Then, this process was iterated to the next most significant peak and so on until all peaks were either kept or removed due to direct overlap with a more significant peak. Differentially accessible peaks were identified using the getMarkerFeatures() function from the ArchR package using a Wilcoxon rank sum test and accounting for bias introduced by TSSEnrichment and log10(nFrags). Peaks with FDR ≤ 0.1 and log2 fold change greater than or equal to 0.25 were considered sex or state specific. Known transcription factor binding motifs present in the CIS-BP database were assigned to differentially accessible genomic regions and motif enrichment was evaluated using a hypergeometic test implemented by the peakAnnoEnrichment() function.

Motifs were then classified into families as defined by TFClass(*101*). The top scoring motifs from each family and across all possible comparisons were then compared for overall enrichment. Motif footprints were calculated by combining all peaks harboring a given motif in aggregate and accounting for Tn5 insertion bias using the getFootprints() function. ChromVAR scores were obtained by running the addDeviationsMatrix() function.

#### DEG analysis

We performed a differential gene expression analysis between samples of different sex and state using a pseudo-bulk approach. We aggregated gene expression values across cell clusters using the Seurat function AggregateExpression() to generate a pseudo-bulk count matrix. After removing genes with zero counts, we defined a design matrix indicating the condition for each sample. Using DESeq2 R library, we constructed a DESeq2 object with DESeqDataSetMatrix() function and performed differential expression analysis with DESeq function with a two-tailed Wald test. We identified significantly differentially expressed genes with an FDR-adjusted p-value inferior to 0.05 and an absolute log2 fold-change value superior to 0.5. We performed GO term enrichment analysis using the R package enrichR. We selected all the GO terms using a one-sided Fisher exact test with an FDR adjusted p-value inferior to 0.05 and visualized the associated adjusted p-value and enrichR combined scores.

#### Random forest classification

To identify which cell types have a notable change in expression between sex and physiological state, we used the R package Augur v.1.0.3 which performs cross-validated random-forest classifier analysis of single-cell datasets. First, genes located on the X or Y chromosome, immediate early genes, as well as mitochondrial and ribosomal genes were removed. Then we applied Augur (var_quantile=0.9, subsample_size=20) to generate area under the curve scores for all possible pair-wise combinations for each cell type.

### Brain slice preparations

Acute slice electrophysiology was performed in adult transgenic Gal:Flp+/-;Calcr:Cre+/-mice, injected with a Cre- and Flp-dependent mScarlet AAV (137136-AAV8) to mark the MPOAGal+Calcr+ neuron population. Acute slice calcium imaging recordings were performed in adult transgenic Brs3:Cre+/-;VgluT2:Flp+/-;Ai195+/- mice to record from the AvPeBrs3+Vglut2+ neuron population. Recordings were made from littermate cohorts of virgin males and virgin females, or siblings mated to become fathers and mothers. Fathers and mothers were recorded from during post-partum day 3-6 and remained with pups until acute slices were made. Estrus cycle was determined for virgin females via vaginal smear on the day of recording. All animals were screened for pup retrieval behavior the day before recording. Only animals exhibiting typical retrieval assay behavior were included in analysis (i.e. attack or ignore for virgin males, retrieve or ignore for virgin females, and retrieve for fathers, and mothers).

#### Solutions

The cutting solution was comprised of 92mM NMDG, 30mM NaHCO3, 25mM Glucose, 20mM HEPES, 2mM Thiourea, 2.5mM KCl, 1.25mM NaH2PO4, 5mM Na-ascorbate, 3mM Na-pyruvate, 0.5mM CaCl2, 10mM MgSO4. Solution was adjusted to pH 7.3-7.4 and 306 mOsm.

The holding solution was comprised of 92mM NaCl, 30mM NaHCO3, 25mM Glucose, 20mM HEPES, 2mM Thiourea, 2.5mM KCl, 1.25mM NaH2PO4, 5mM Na-ascorbate, 3mM Na-pyruvate, 2mM CaCl2, 2mM MgSO4. Solution was adjusted to pH 7.3-7.4 and 306 mOsm. The ACSF was comprised of 119mM NaCl, 24mM NaHCO3, 12.5mM Glucose, 2.5mM KCl, 1.25mM NaH2PO4, 2mM CaCl2, 2mM MgSO4. Solution was adjusted to pH 7.3-7.4 and 306 mOsm. The current-clamp internal solution was comprised of 120mM K-gluconate, 10mM Na-gluconate, 10mM HEPES, 10mM Na-phosphocreatine, 4mM NaCl, 4mM Mg-ATP, 2mM Na2-ATP, 0.3mM Na3-GTP, 4mM Biocytin. Solution was adjusted to pH 7.35 and 297 mOsm.

#### Acute brain slices

Acute slices were prepared as described in Ting, et al.(*102*), Briefly, adult mice were transcardially perfused with ice-cold NMDG-based cutting solution. Brains were extracted in fresh ice-cold cutting solution and mounted for coronal sectioning (300µm thick) in continuously oxygenated ice-cold cutting solution on a Campden 7000smz-2 vibratome. Following sectioning, slices were moved to continuously oxygenated warmed cutting solution in a 34°C water bath and incubated for 11 minutes. Slices were then moved to continuously oxygenated holding solution at room temperature to recover for 1 hour.

#### In vitro recordings

Whole-cell current clamp recordings were made using borosilicate glass patch pipettes (3-8 MΩ) filled with current-clamp internal solution. Slices were maintained under constant perfusion with warmed oxygenated ACSF at 32°C supplemented with synaptic blockers (20µM Bicuculline (Tocris, Cat # 14340), 50µM AP5 (AbCam, Cat # ab120003), 10µM CNQX (Sigma, Cat # C239)) for the duration of recording. Slices were visualized on an Axio Examiner (Zeiss, Oberkochen, Germany). Recordings were obtained with a Multiclamp 700B amplifier (Molecular Devices, Palo Alto, CA) and 1550B4 Digidata digitizer (Molecular Devices), and physiological data were collected via pClamp11 (Molecular Devices).

Frequency-current curves (F-I curves) were measured in response to increasing 10pA current steps, starting at −50pA, until 200pA or the cell reached depolarization block. Rheobase measurements were made by applying successive 1pA current steps, starting at 0pA, and the current at which the cell fired a single action potential was measured. All electrophysiological measurements were repeated at least twice for each cell and an average of these measurements is reported. Analysis of electrophysiological recordings was performed with Clampfit11 (Molecular Devices) and Axograph (Axograph) software.

#### Calcium imaging recordings

Following 1 hour recovery, acute slices from Brs3:Cre+/-;VgluT2:Flp+/-;Ai195+/-transgenic mice were incubated on the imaging rig under constant perfusion with warmed oxygenated ACSF at 32°C and with the objective lowered into the bath for an additional 1 hour to prevent focusing aberrations due to thermal drift. Calcium signals from GCaMP7s+ cells were recorded on an Axio Examiner (Zeiss) microscope with Zen Blue 3.3 software (Zeiss). A 20x W Plan-Apo objective was used to image cells at 5Hz, 75ms exposure time, 5% LED power and 30% intensity. GCaMP7s was excited using the Colibri light engine (Zeiss). Following a 2-minute baseline, ACSF supplemented with the oxytocin receptor-specific agonist 2µM [Thr4,Gly7]-oxytocin, aka TGOT (Bachem, Cat # 4013837.0005) was applied for 5 minutes to measure response to oxytocin receptor activation.

For analysis, videos were first motion corrected and regions of interest identified through the CaImAn software package (*103*) For some videos it was necessary to manually annotate regions of interest using imagej. After correction for florescence bleaching, relative changes in fluorescence (ΔF/F) were calculated for each region where F is the mean baseline fluorescence intensity) unless otherwise stated.

#### Statistics and reproducibility

Data were processed and analyzed using a combination of R and Python codes. Sample sizes were chosen based on common practice in single-nucleus sequencing, electrophysiology, and animal behavior experiments. Individual data points were plotted wherever possible. Boxplots represent the median, first and third quartiles (hinges) and 1.5× interquartile range (whiskers). Outliers are shown wherever individual data points are not plotted. Behavioral experiments were analyzed using Kaplan-Meier survival analysis with log rank test, and all other data were analyzed using two-tailed non-parametric tests unless indicated otherwise. *P < 0.05, **P < 0.01, ***P < 0.001, and ****P < 0.0001. Statistical details are given in the respective figure legends.

## Supplemental figure legends

**Fig. S1.**
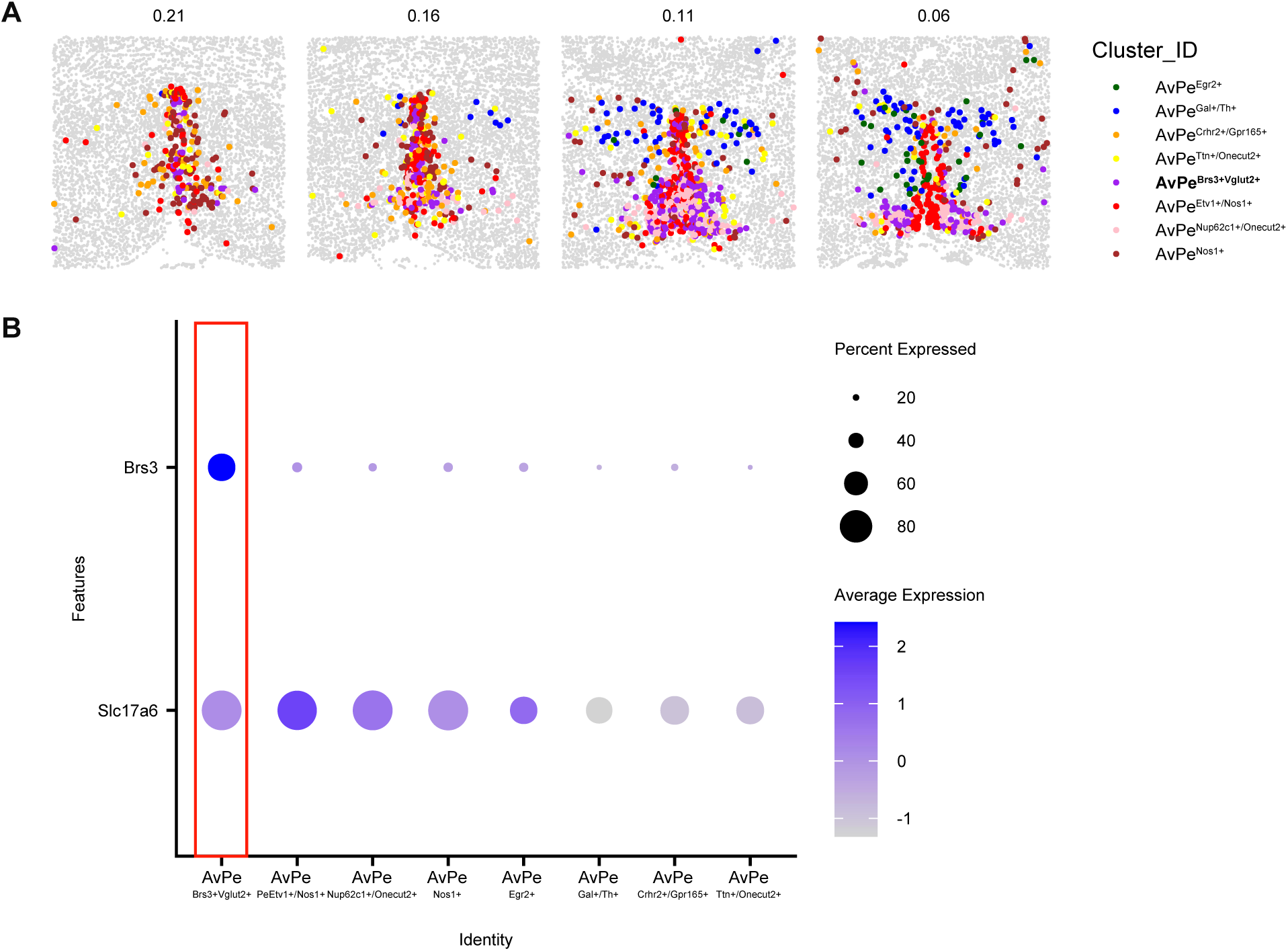
Brs3 and Slc17a6 serve as molecular markers for AvPe^Brs3+Vglut2+^ neurons. (**A**) MERFISH images of molecularly defined AvPe cell types (*9*)(**B**) Gene expression of AvPe cell types indicating expression profiles that are present in the desired population.

**Fig. S2.**
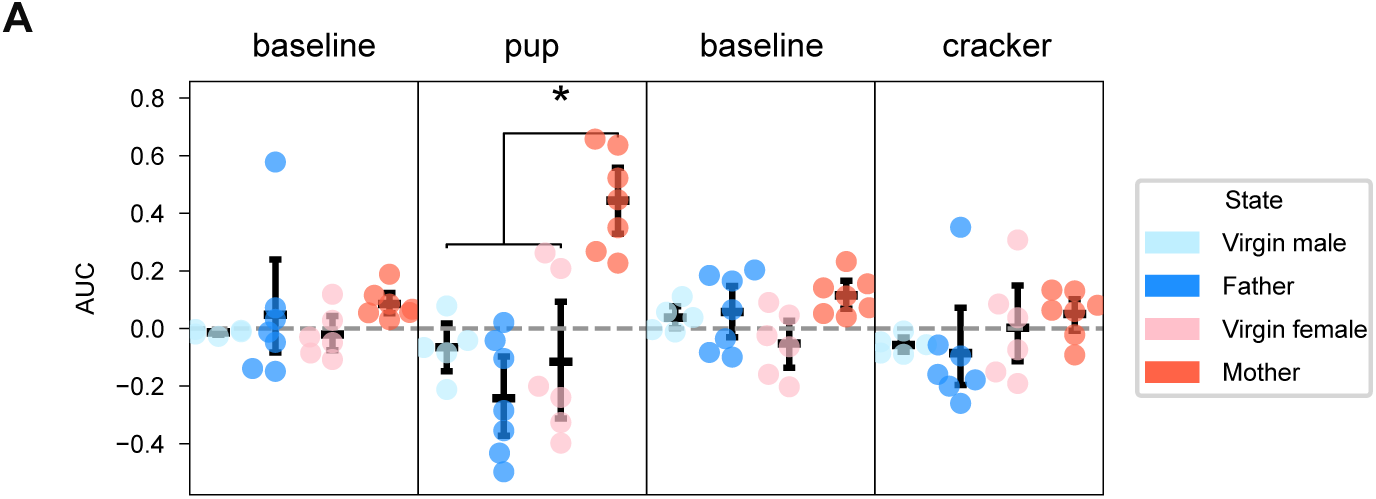
AvPe^Brs3+Vglut2+^ neurons are active during pup exposure. (**A**) Cumulative area under the curve of GCaMP7s activity during the duration of assay exposure for all animals.

**Fig. S3.**
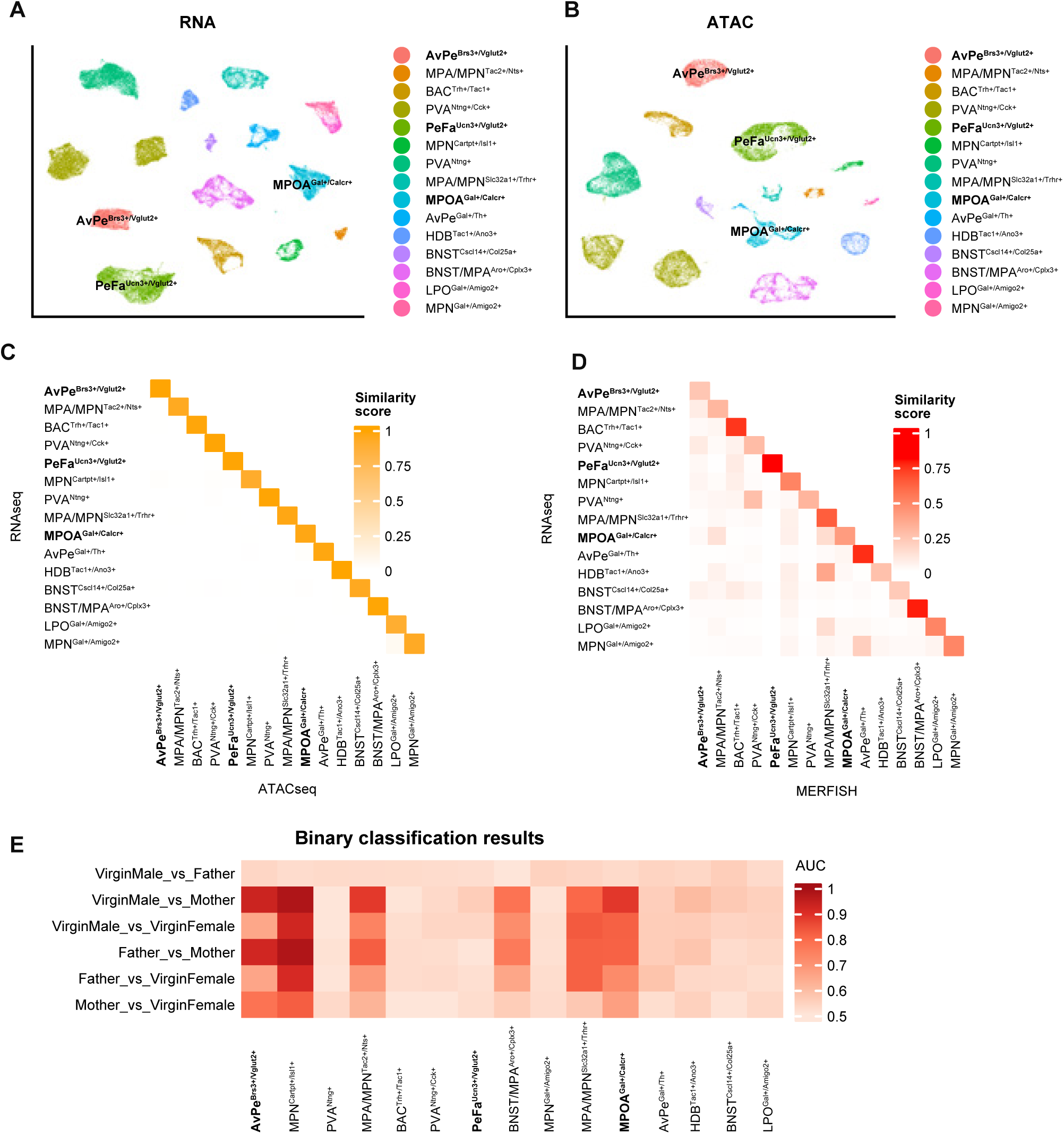
RNA- and ATAC-seq data map to previous cell types and reveal unique expression patterns. (**A-B**) UMAP plot highlighting the distinct nature of the identified cell types for both molecular modalities. (**C**) Canonical correlation score calculated from each cluster compared to all others for RNA- and ATAC-seq modalities. (**D**) Same as in (**C**) but comparing the RNA-seq dataset to the previously published MERFISH dataset from *Moffitt et al.* (**E**) Area under the curve score for cell type-specific random forest classification results of all pair-wise decoding combinations.

**Fig. S4.**
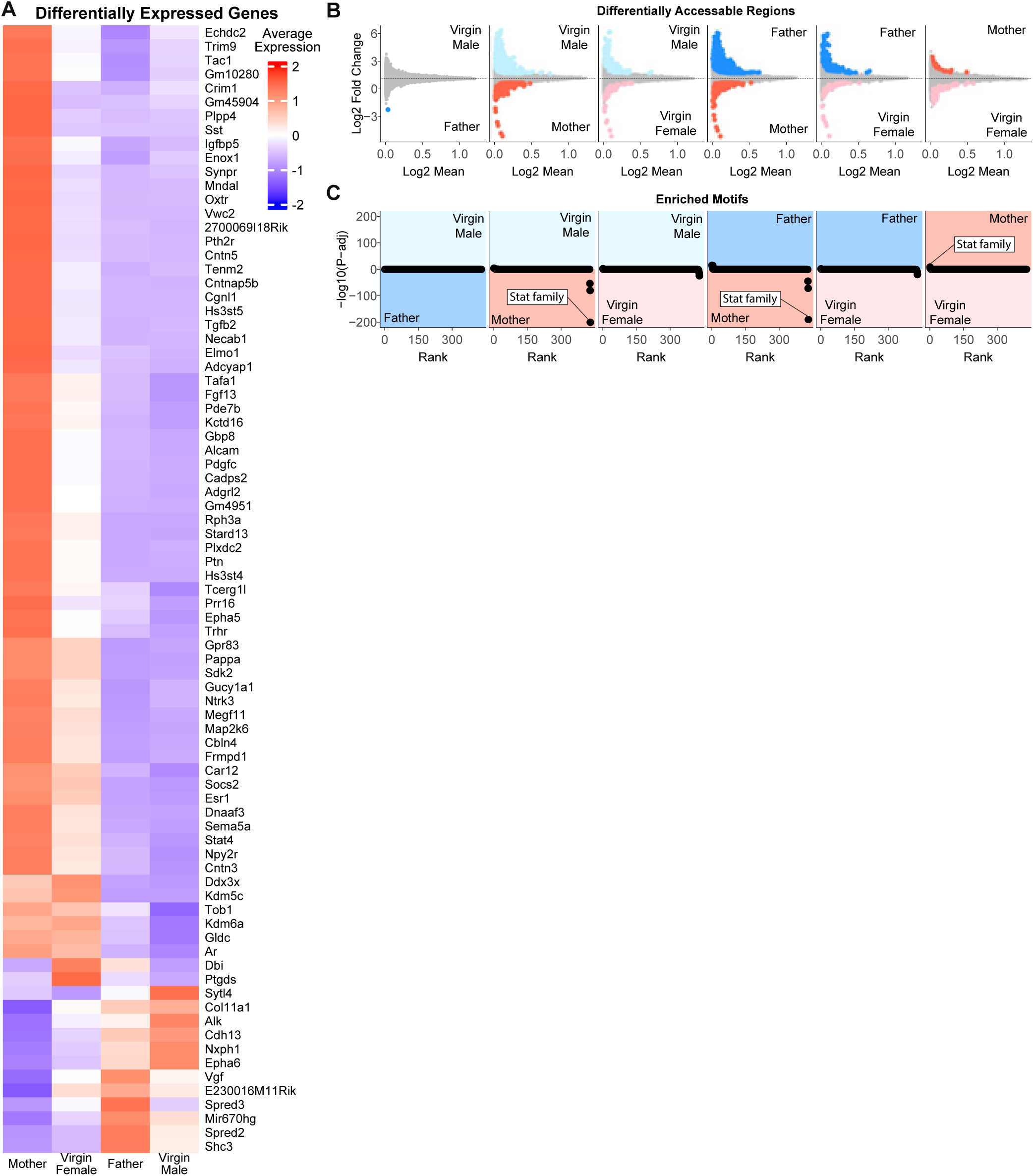
AvPe^Brs3+Vglut2+^ neurons show unique molecular profile in mothers. (**A**) All 81 differentially expressed genes between animal sex and state hierarchically clustered. (**B**) Pair-wise log2Fold change plots of differentially accessible regions for all combinations of samples. Color represents significant differences in accessibility. (**C**) Pair-wise -log10 plots of transcription factor motifs enriched in accessible peaks for all combinations of samples.

**Fig. S5.**
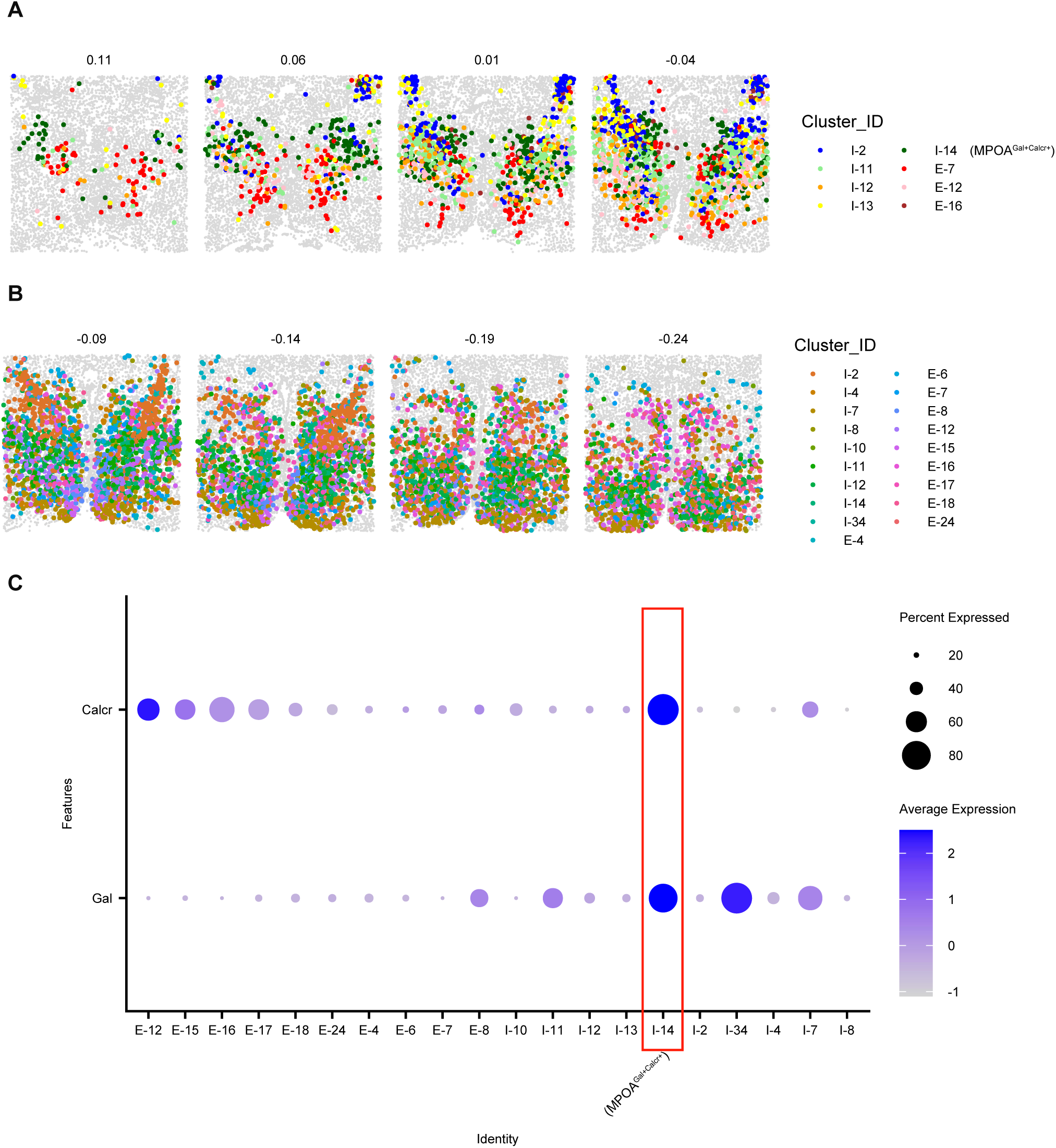
Gal and Calcr serve as molecular markers for MPOA^Gal+Calcr+^ neurons. (**A**) MERFISH images of defined cell types from Moffit et al located in the StHy. (**B**) MERFISH images of defined cell types from Moffit et al located in the MPN. (**C**) Gene expression of StHy and MPN localized cell types indicating expression profiles that are present in the desired population.

**Fig. S6.**
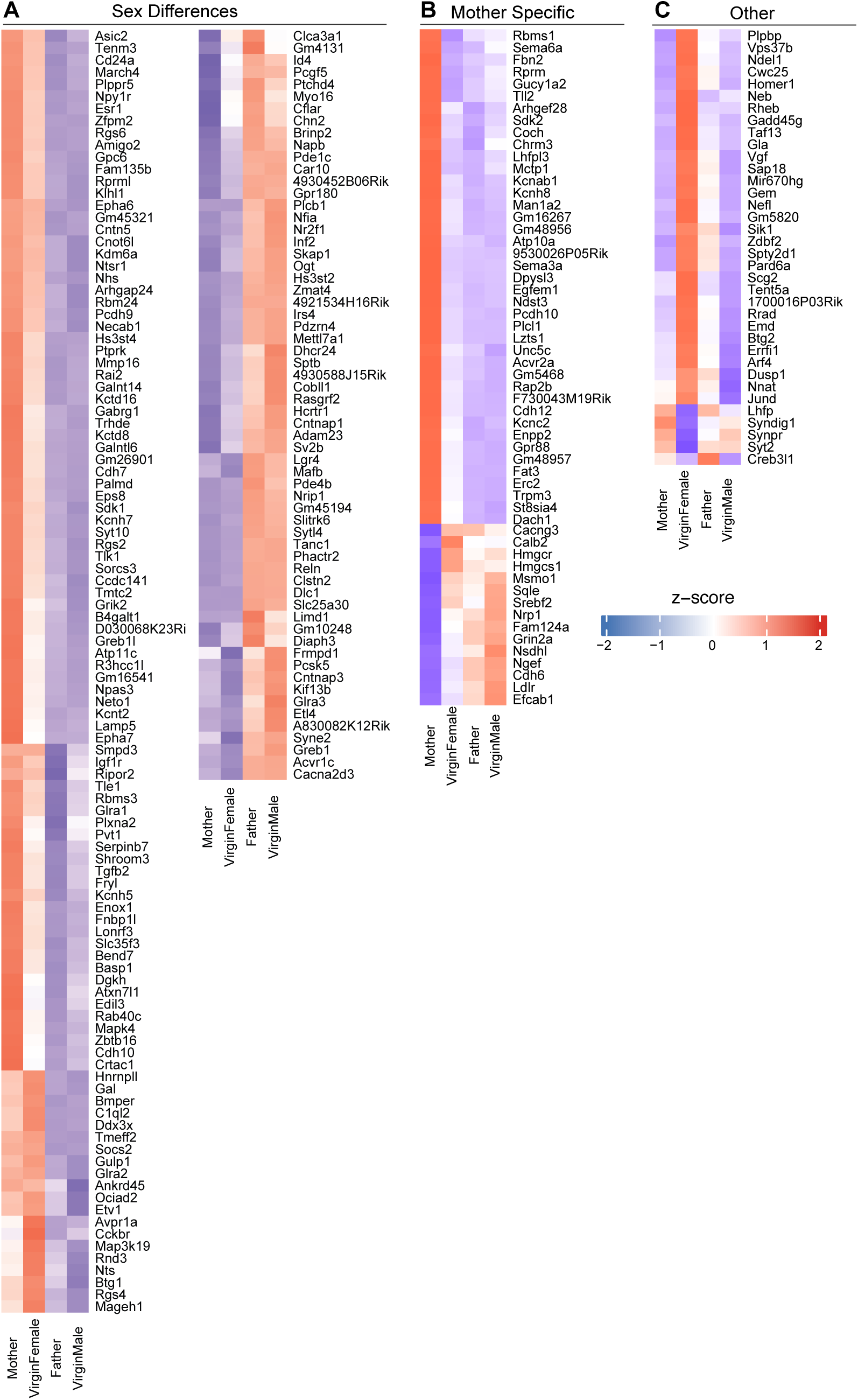
MPOA^Gal+Calcr+^ neurons show complex differences in gene expression patterns. (**A**) Genes that show sex differences in expression. (**B**) Genes showing expression changes in mothers compared to all other comparisons. (**C**) Remaining differentially expressed genes showing additional expression patterns.

**Fig. S7.**
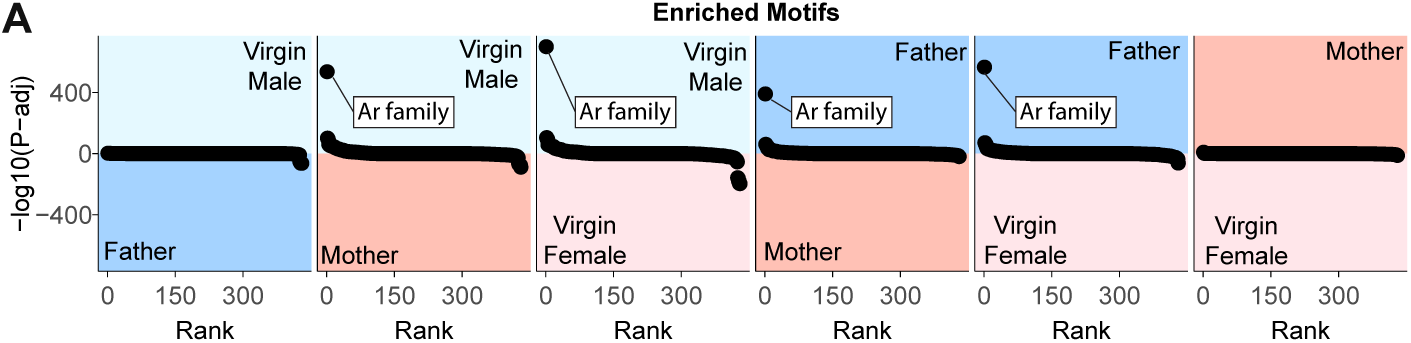
MPOA^Gal+Calcr+^ neurons show complex differences in chromatin accessibility and motif enrichment. (**A**) Pair-wise -log10 plots of transcription factor motifs enriched in accessible peaks for all combinations of samples.

**Fig. S8.**
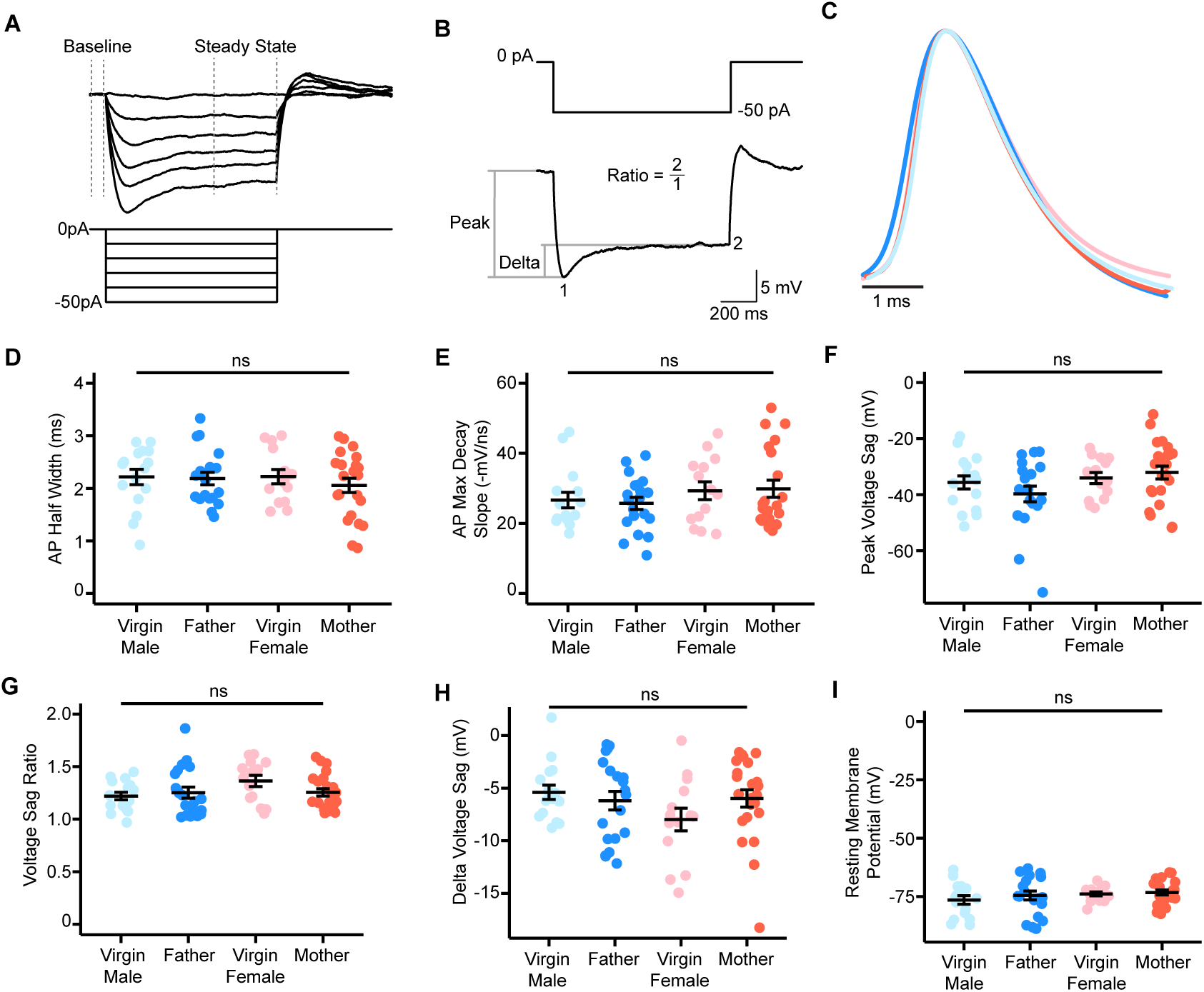
Electrophysiological features of MPOA^Gal+Calcr+^ neurons. (**A**) Example trace demonstrating how input resistance was calculated. Measured injected current was plotted versus measured average steady state voltage (minus baseline voltage) for each current step. Input resistance was calculated as the slope of a line fitted to these data. (**B**) Example trace demonstrating how voltage sag was calculated. (**C**) Scaled average action potential traces across virgin males, fathers, virgin females, and mothers. (**D**) Action potential half width (Two-way ANOVA, no significant effects, father n = 19, mother n = 22). (**E**) Action potential maximum decay slope (Two-way ANOVA, no significant effects, virgin male n = 15, mother n = 21). (**F**) Peak voltage sag of cells (Two-way ANOVA, no significant effects). (**G**) Voltage sag ratio of cells (Two-way ANOVA, no significant effects). (**H**) Resting membrane potential of cells (adjusted for liquid-liquid junction potential, Two-way ANOVA, no significant effects). (**I**) Delta voltage sag of cells (Two-way ANOVA, no significant effects). (**J**) Number of animals performing each behavior that were used for recording. Error bars are mean ± SEM, ns = non-significant, * = *p* < .05. Unless otherwise noted, virgin male n = 16, father n = 20, virgin female n = 14, mother n = 23.

